# Formin-2 drives intracellular polymerisation of actin filaments enabling correct segregation of apicoplasts in *Plasmodium falciparum* and *Toxoplasma gondii*

**DOI:** 10.1101/488528

**Authors:** Johannes Felix Stortz, Mirko Singer, Jonathan M Wilkes, Markus Meissner, Sujaan Das

## Abstract

Pathogenic obligate-intracellular apicomplexan parasites possess an essential chloroplast-like organelle called the apicoplast that undergoes division and segregation during replication. Parasite actin is essential during intracellular development, implicated in vesicular transport, parasite replication and apicoplast inheritance. However, the inability to visualise live actin dynamics in apicomplexan parasites limited functional characterisation of both filamentous-actin (F-actin) and actin regulatory factors. Apicomplexans possess at least two distinct formins, Formin-1 and Formin-2, predicted to serve as actin-nucleating factors, and previously implicated in regulating gliding motility and host cell invasion. Here, we expressed chromobodies and validated them as F-actin-binding sensors in *Plasmodium falciparum* and characterised the *in vivo* dynamics of the F-actin network. The F-actin network could be modulated chemically and disrupted by conditionally deleting the *actin-1* gene. In a comparative approach, we demonstrate that Formin-2 is closely associated with apicoplasts and with the F-actin network in *P. falciparum* and *Toxoplasma gondii*. Consequently, disruption of Formin-2 resulted not only in an apicoplast segregation defect, but also in complete abrogation of F-actin dynamics in intracellular parasites. Together, our results strongly indicate that Formin-2-mediated filament formation is the common primary mechanism for F-actin nucleation during apicomplexan intracellular growth effecting apicoplast segregation.

## Introduction

The phylum Apicomplexa includes a variety of obligate intracellular parasites, which invade into and replicate inside mammalian cells, causing immense disease burden in humans and in commercially important livestock. One of its notorious members, the malaria parasite *Plasmodium falciparum*, is a major health concern in developing nations, causing ~500,000 deaths annually (White, Pukrittayakamee et al. 2014). Another member, *Toxoplasma* is a highly successful parasite infecting almost a third of the global human population and can be fatal in immunocompromised patients (Torgerson and Mastroiacovo 2013). There is limited success in the development of vaccines against these parasites and the current drugs are associated with drug resistance, making it crucial to investigate novel biological targets.

Actin is one of the most abundant proteins in eukaryotic cells. Due to its ability to form polymers, this cytoskeletal protein is involved in numerous processes such as cell motility, cytokinesis, organellar and vesicular transport, secretion and endocytosis (Svitkina 2018). Actins encoded by apicomplexan parasites are highly divergent compared to canonical actins from other eukaryotes (Douglas, Nandekar et al. 2018). *In vitro*, apicomplexan actins form only short, unstable polymers due to different polymerisation kinetics, caused by variation of certain key amino acids, otherwise conserved in metazoans (Kumpula and Kursula 2015). However, until recently an analysis of F-actin localisation and dynamics in apicomplexan parasites was hindered by the unavailability of F-actin sensors (Tardieux 2017), a limitation recently overcome by the expression of F-actin binding chromobodies in *T.gondii* (Periz, Whitelaw et al. 2017). Intriguingly, in this parasite, F-actin can form an extensive intra-vacuolar network that appears to be involved in material exchange and synchronisation of parasite division (Periz, Whitelaw et al. 2017).

Until recently, studies on apicomplexan F-actin focused on its critical role during host cell invasion and gliding motility (Soldati, Foth et al. 2004, Baum, Gilberger et al. 2008), where it is believed to provide the force for both processes (Frenal, Dubremetz et al. 2017). However, recent studies using conditional mutants for *actin-1* in two apicomplexans, *P. falciparum* and *T. gondii* highlight additional critical roles of F-actin during intracellular parasite development (Das, Lemgruber et al. 2017, Periz, Whitelaw et al. 2017, Whitelaw, Latorre-Barragan et al. 2017). Intriguingly, some functions, such as inheritance of the chloroplast-like organelle, the apicoplast, appears to be conserved (Andenmatten, Egarter et al. 2013, Egarter, Andenmatten et al. 2014, Das, Lemgruber et al. 2017, Whitelaw, Latorre-Barragan et al. 2017), while differences for the dependency of F-actin can be observed for other critical steps of the asexual life cycle. For example, host cell invasion is possible without *actin-1* (albeit at highly reduced levels) in case of *T. gondii* (Andenmatten, Egarter et al. 2013, Egarter, Andenmatten et al. 2014, Whitelaw, Latorre-Barragan et al. 2017), while it is completely blocked in case of *P. falciparum* (Das, Lemgruber et al. 2017). In contrast, *P. falciparum* does not require actin dynamics for egress from the host cell (Das, Lemgruber et al. 2017, Perrin, Collins et al. 2018), while it is essential for *T.gondii*.

Of the two actin genes present in *P. falciparum* (Gardner, Hall et al. 2002), only *actin-1* (*pfact1*) is expressed in all life-cycle stages and is the only actin expressed during asexual replicative stages, whereas *actin-2* expression is confined to the sexual gametocyte and insect stages (Vahokoski, Bhargav et al. 2014). *P. falciparum* undergoes a 48h asexual replicative cycle in the intermediate human host where it invades into, grows and replicates within erythrocytes, causing all clinical manifestations of the disease. After invasion, the merozoite form of the parasite (similar to *T. gondii* tachyzoites) establishes itself within a parasitophorous vacuole (PV), loses its ovoid shape to become amoeboid and feeds on host haemoglobin creating a food vacuole (FV) where haem is detoxified (Gruring, Heiber et al. 2011). The parasite then replicates by a process best described as internal budding, where daughter parasites develop within the mother (Francia and Striepen 2014). In the case of *T. gondii*, only two daughters are formed at a time in a process called endodyogeny. In contrast, malaria replication within the erythrocyte, termed schizogony, results in the formation of 16-32 merozoites at once. Towards the end of a replicative cycle the parasite *de novo* forms its invasion-related organelles: the inner membrane complex (IMC), micronemes and rhoptries. In contrast, parasite mitochondria and the apicoplast undergo growth and division and are trafficked into each daughter cell (Bannister, Hopkins et al. 2000). Although endodyogeny and schizogony appear very different, it is believed that both processes are very similar and use conserved molecular machinery. Indeed, independent studies identified the same factors to be critical for both replicative modes (Francia and Striepen, 2014).

Despite this, it could be assumed that differences, especially with respect to vesicular transport processes such as endocytosis and intravacuolar parasite communication, exist to adapt to different replication modes. This puts F-actin in the spotlight, since it plays a central role in these processes, as is the case in other eukaryotes (Svitkina 2018). We recently characterised a conditional mutant of PfACT1 and observed that in good agreement with the function of actin in *T. gondii* (Andenmatten, Egarter et al. 2013), inheritance of the apicoplast is compromised during schizogony (Das, Lemgruber et al. 2017). While the phenotypic analysis of conditional mutants is useful to identify conserved and unique functions of F-actin in apicomplexans, the inability to visualise F-actin in these parasites led to models, sometimes conflicting with each other and with the canonical behaviour of F-actin in other eukaryotes.

Common actin-labelling probes such as Phalloidin do not label apicomplexan actin and LifeAct could not be successfully expressed in these parasites (Periz, Whitelaw et al. 2017). Recently, actin-binding single-domain nanobodies tagged to fluorescent probes, called chromobodies were successfully expressed in *T. gondii* and shown to have minimal effect on actin dynamics (Periz, Whitelaw et al. 2017), as also demonstrated in other eukaryotic cells (Rocchetti, Hawes et al. 2014, Panza, Maier et al. 2015, Melak, Plessner et al. 2017).

Here we adapted this technology to *P. falciparum* and demonstrate for the first time the localisation, dynamics and role of F-actin dynamics for parasite development in asexual stages. Interestingly, we find F-actin closely associated with the apicoplast throughout intracellular growth, leading to the question of which actin regulatory proteins are involved in this process. Most actin nucleation proteins such as the Arp2/3 complex and the WAVE/WASP complex, and actin cross-linkers such as α-actinin and fimbrin are missing in apicomplexans (Baum, Papenfuss et al. 2006, Schuler and Matuschewski 2006). Two conserved nucleators found in *P. falciparum* are the formins, Formin-1 (PfFRM1) and Formin-2 (PfFRM2) which localise to distinct compartments in the cell (Baum, Tonkin et al. 2008). Orthologs of both formins have been implicated in host cell invasion in *T. gondii* (Daher, Plattner et al. 2010), with *T. gondii* Formin-2 (TgFRM2) also being implicated in apicoplast maintenance (Jacot, Daher et al. 2013) - leading to inconsistencies in reports and questions whether the two formins have conserved or divergent functions in both parasites.

Here we reanalysed the role of Formin-2 in *P. falciparum* and *T. gondii* and, in contrast to previous reports, demonstrate that it localises adjacent to apicoplasts in both parasites. Conditional disruption of Formin-2 not only results in a complete abrogation of actin dynamics in *P. falciparum* and *T. gondii*, it also leads to loss of the apicoplast. Together our study highlights a highly conserved role of Formin-2 in the intracellular development of apicomplexan parasites. Importantly, apicoplast loss appears to be not the only critical phenotype caused by Formin-2 depletion, since in *P. falciparum* the loss of fitness due to the deletion of PfFRM2 cannot be complemented by addition of isopentenyl pyrophosphate (IPP), the only essential metabolite produced by apicoplasts (Yeh and DeRisi 2011), suggesting other critical roles of Formin2-mediated actin nucleation in these parasites.

## Results

### 1. Cellular expression of chromobodies in *P. falciparum* enables the visualisation of an actin network throughout the asexual development of *P. falciparum* and in gametocytes

#### a. Chromobodies label F-actin structures in *P. falciparum* asexual stages and in gametocytes

Chromobodies were expressed under the *heat shock protein 86 (hsp86)* promoter (Crabb and Cowman 1996) to obtain expression throughout the 48h asexual life cycle. We succeeded in generating parasites stably expressing chromobodies tagged either with the emerald tag (CB-EME) or the halo tag (CB-HALO), indicating that the expression of these constructs does not have a major deleterious impact on the fitness of *P. falciparum* (**Fig. 1A**), as previously reported for *Toxoplasma* (Periz, Whitelaw et al. 2017) and other eukaryotes. The halo tag allowed visualisation of F-actin in live parasites by use of the red ligand Halo-TMR. Dynamic filamentous structures were evident in both CB-EME and CB-HALO expressing parasites (**Fig 1B**) throughout the 48 hour life cycle (**Videos V1, V2, V3 and V4**). These structures could be completely disrupted by adding the F-actin destabilising drug cytochalasin-D (**Fig. 1B** and **Video V3**), demonstrating that chromobodies bind F-actin structures in *P. falciparum*. However, while both CB-versions labelled similar structures, we found that expression of CB-EME resulted in a better signal-to-noise ratio, probably because no permeable, fluorescent ligand needs to be added (**Fig.1B**). Therefore, for the rest of this study results for parasites expressing CB-EME are presented. Some F-actin structures were highly dynamic, changing within a time-scales of seconds, while other structures appeared stable over tens of seconds (**Video V3**). For better visualisation of F-actin structures, we used super-resolution microscopy (SR-SIM) which enabled us to observe a complex F-actin network in these parasites (**Figs. 1C** and **S1**), similar to a network observed for *T.gondii* (Periz, Whitelaw et al. 2017). Since CB-EME was expressed from an episome, we quantified mean intensity of fluorescence in different cells and found it to be within a narrow range (**Fig. S1B**). Interestingly, F-actin was most prominent around the FV of the parasite (**Figs. 1C** and **S1**). When we co-stained the chromobody-labelled network with an antibody which specifically recognises parasite actin, we observed a rather cytosolic distribution with the antibody, although similar structures as seen with chromobody were apparent. This indicates that the antibody recognises monomeric globular actin (G-actin) along with F-actin, while the chromobody recognises primarily filamentous structures (**Fig. 1C** and **Fig S1**). In contrast to asexual parasites, gametocytes express both PfACT1 and actin-2, and exhibit F-actin staining along the length of the parasite and at the tips (Hliscs, Millet et al. 2015). Upon expression of CB-EME, gametocytes showed intense dynamic F-actin structures at their tips and running along the whole body of the cell changing in seconds (**Fig. 1D** and **Video V5**). Importantly, this dynamic network appears very similar to the one reported by Hlisc et al., 2015, which has been shown to lie beneath the IMC of the gametocyte. It is important to note that chromobodies do not distinguish between PfACT1 and actin-2 and therefore the observed filaments could be built from both proteins. Together, our data show that expression of chromobodies does not cause significant phenotypic effects and allow reliable labelling of the F-actin cytoskeleton. We also confirm previous data obtained with antibodies directed against F-actin and show that during the gametocyte stage, F-actin forms a dynamic and extensive network that passes through the whole cell and is enriched at the tips of the parasite.

**Figure 1.**
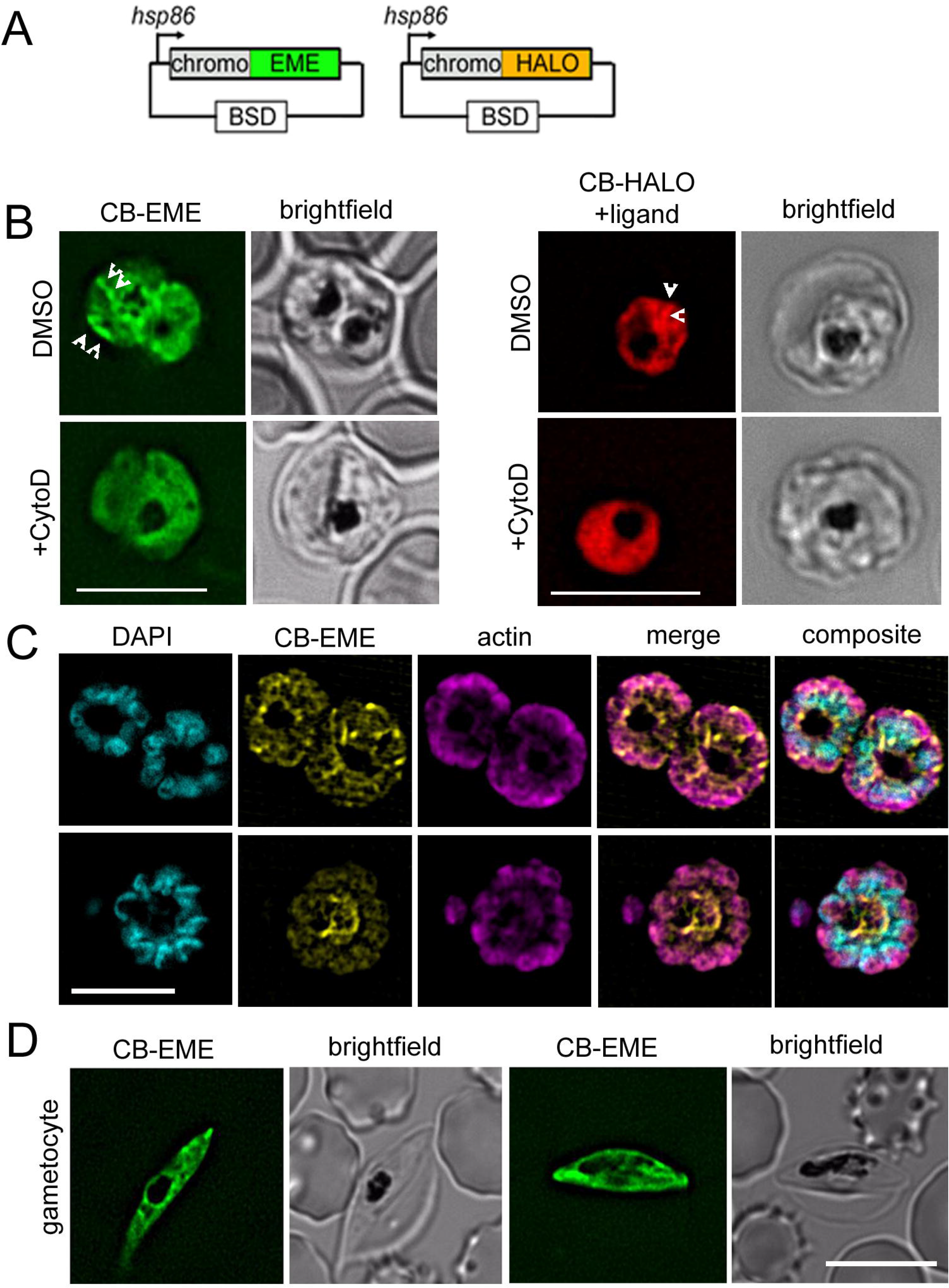
Chromobody constructs with different epitope tags label F-actin structures throughout the *P. falciparum* lifecycle. Chromobody constructs used in this study under the *hsp86* promoter with an emerald tag (CB-EME) and halo tag (CB-HALO). Blasticidin-S-deaminase (BSD) confers resistance to blasticidin. **B.** CB-EME and CB-HALO label actin filaments (DMSO), which disappear upon cytochalasin-D treatment (+CytoD). White arrows denote examples of F-actin structures. Also see **Video V3**. **C.** Super resolution imaging reveals an actin network in *P. falciparum* (CBEME), which stains partially similar to an actin antibody (actin). DAPI labels nuclei. See also **Fig. S1**. **D.** The F-actin network and dynamics can be visualised in gametocytes (see also **Video V5**). Brightfield images provided in greyscale alongside. Scale bar 5μm.

#### b. Highly dynamic F-actin-rich filopodia-like structures extend outward from the periphery of growing parasites

After erythrocyte invasion, the parasite immediately loses its ovoid zoite structure and becomes an amoeboid ring-stage parasite. These young parasites are highly dynamic and can switch between various shapes forming multi-lobed structures, possibly mediated by their cytoskeletal networks (Gruring, Heiber et al. 2011). On observing chromobody-expressing parasites during ring and early trophozoite stages we noted F-actin rich projections at the periphery of ~60% parasites (n=50) (**Fig. 2A, B** and **Video V2**). These F-actin projections are highly dynamic, changing in orders of seconds. Upon treatment with the F-actin depolymerising drug cytochalasin-D the peripheral dynamic F-actin protrusions disappeared (**Fig. 2A lower panel**), but the multilobed structures of the parasite were not disrupted (**Video V6**). The F-actin stabilising drug jasplakinolide also disrupted these filopodia-like extensions and resulted in formation of thick filaments (**Fig 4A lower panel**), implying the requirement of dynamic regulation of F-actin for maintenance of these projections. The physiological relevance of these filopodia is unclear at this point and further research into this phenomenon is required.

**Figure 2.**
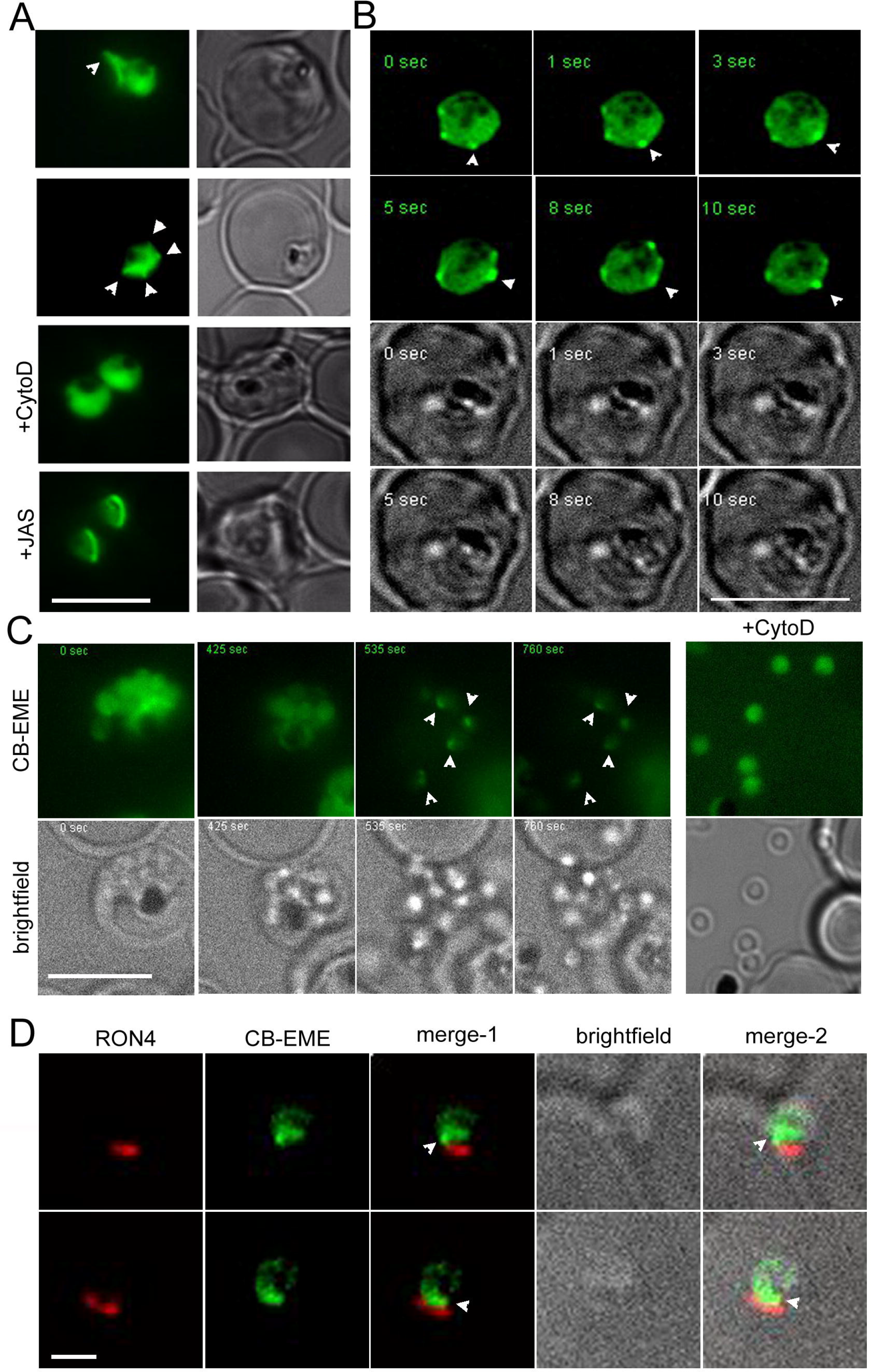
Rapid actin dynamics are visible during intracellular growth, egress and invasion. **A.** Filopodia-like protrusions from the parasite cell body extend into the host cell cytosol (white arrows), which are disrupted by cytochalasin-D (+CytoD) and jasplakinolide (+JAS). **B.** Time lapse images show rapid changes in filopodia-like protrusions (white arrows). See also **Video V2**. CB-EME visible in the green channel, and brightfield images have been provided below in greyscale. **C.** Time lapse images show actin polymerising at a polar end of the merozoite, post egress (CB-EME). Corresponding brightfield images have been presented. Cytochalasin-D treatment prohibits the polar polymerisation of actin (+CytoD). **D.** IFA of invading merozoites with the junction marker RON4 shows CB-EME staining close to the RON4 stain, implying that F-actin polymerises at the apical end prior to invasion.

#### c. Apical polymerisation of F-actin in merozoites following egress

Next, we wished to analyse the fate of the observed F-actin network upon parasite egress. We synchronised parasites with a 2-step Percoll and sorbitol treatment and harvested schizonts at 44h post-invasion. Reversible inhibitors of protein kinase G, Compound-1 and −2, stall schizont development at very mature stages without allowing them to undergo egress (Collins, Hackett et al. 2013). We treated highly mature schizonts with Compound-2 to allow them to fully mature without undergoing egress for 4h. Upon washing away Compound-2, the parasites egressed normally with the concomitant appearance of F-actin accumulation at the apical tip of the parasite (**Fig. 2C**, **Video V4**). Cytochalasin-D treatment allowed normal egress of parasites, as previously observed (Video S8 of (Weiss, Gilson et al. 2015)), but completely abrogated F-actin polymerisation at the apical tip (**Fig. 2C**). Finally, we performed IFA on invading merozoites using rhoptry neck protein 4 (RON4) as junctional marker. We verified that F-actin accumulates just behind the RON4 ring (**Fig. 2D**) confirming previous observations (Riglar, Richard et al. 2011, Angrisano, Riglar et al. 2012).

#### d. Chromobody labelled F-actin structures disappear upon disruption of PfACT1

Although *P. falciparum* parasites possess two actin genes *pfact1* and *pfact2*, PfACT1 is the only protein expressed during the asexual life cycle (Vahokoski, Bhargav et al. 2014). In order to confirm that chromobodies label authentic F-actin structures based on polymerisation of PfACT1, we transfected the chromobody constructs CB-EME (**Fig. 3A**) and CB-HALO (**Fig. S2B, C**) into a conditional mutant of PfACT1 (loxPACT1) (Das, Lemgruber et al. 2017) (**Fig 3A**). Upon activation of DiCre with rapamycin, the *pfact1* locus is excised together with loss of PfACT1 protein within 35h (Das, Lemgruber et al. 2017). Upon disruption of *pfact1* in 1h-old ring stages, CB-EME (**Fig. 3B, D, E**, **Video V7**) and CB-HALO (Fig. S2) labelled F-actin structures completely disappeared in late trophozoites and schizonts and closely resembled parasites treated with cytochalasin-D (**Fig. 1B**). As previously reported (Das, Lemgruber et al. 2017), PfACT1-disrupted parasites could not invade new erythrocytes (**Fig. 3C**).

We previously found that apicoplast inheritance depends on PfACT1 (Das, Lemgruber et al. 2017). In order to determine the localisation of F-actin during apicoplast segregation we used deconvolution microscopy on fixed parasites stained with the apicoplast marker CPN60, which revealed a close apposition of apicoplasts with F-actin structures (**Fig 3F** DMSO). Upon disruption of PfACT1, a defect in apicoplast segregation was apparent (**Fig 3F** RAP) recapitulating the phenotype observed previously (Das, Lemgruber et al. 2017). Super-resolution microscopy confirmed that actin filaments labelled by CB-EME are closely placed next to migrating apicoplasts (**Fig S1**, **lower panel**). In contrast, no obvious defects in mitochondria segregation could be detected in PfACT1-disrupted parasites (**Fig. 3E** and **Video V7**) as previously reported (Das, Lemgruber et al. 2017), implying that unlike apicoplasts, mitochondria do not require F-actin for migration into daughter cells. IMC markers GAP45 and MTIP showed normal staining in CB-EME expressing parasites (**Fig S1**).

**Figure 3.**
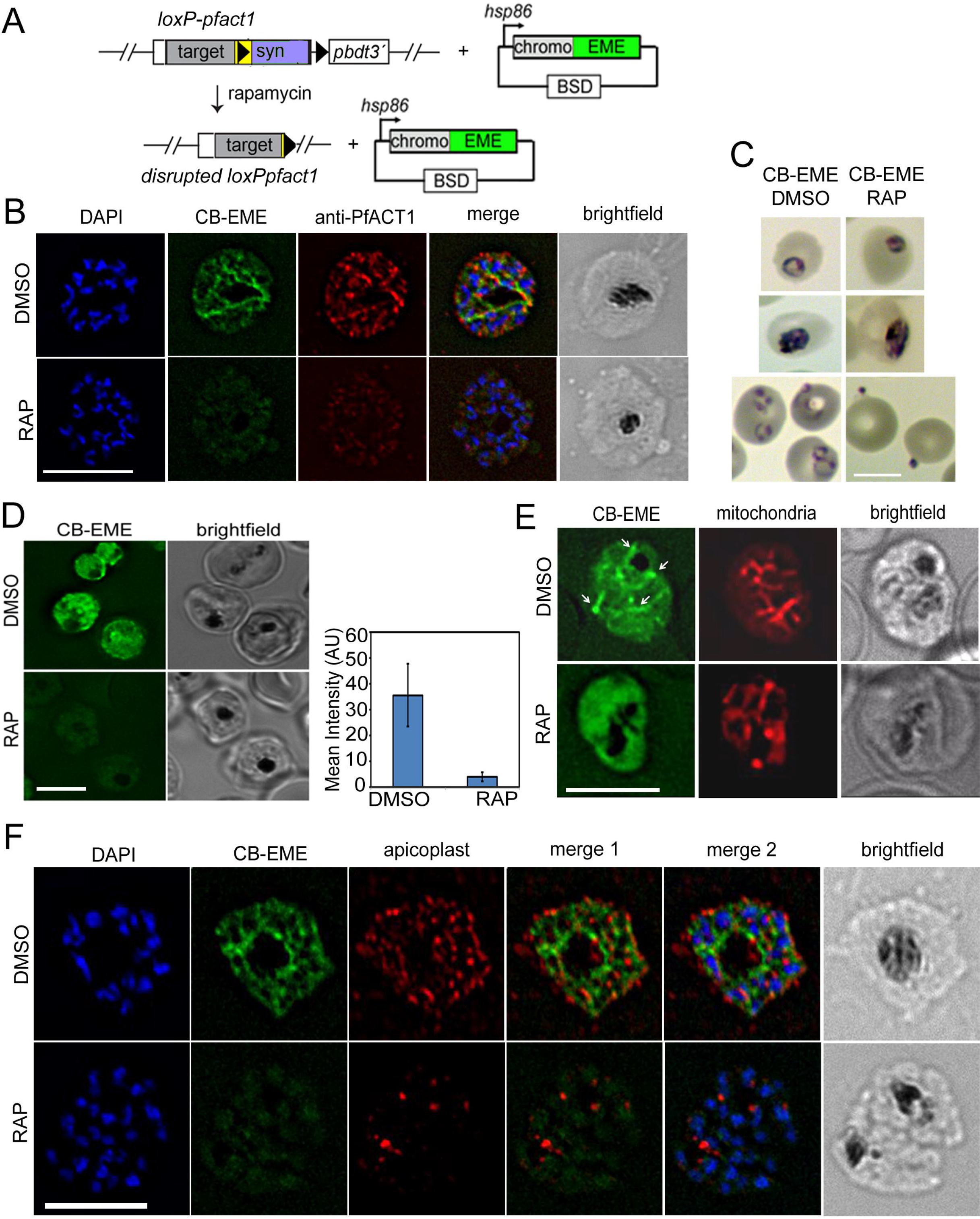
CB-EME staining of the F-actin network disappears upon genetic disruption of PfACT1. **A.** Schematic of transfection of CB-EME into the loxPpfact1 strain and PfACT1 loss upon DiCre-activation using rapamycin. **B.** The F-actin network (CB-EME, green and anti-PfACT1, red DMSO) is lost upon DiCre-mediated disruption of PfACT1 (RAP). DAPI labels nuclei. **C.** Giemsa-stained parasites showing invasion is abrogated in PfACT1 KOs. **D.** Live microscopy of CB-EME expressing parasites and the loss of fluorescence intensity upon RAP-treatment. Right panel shows quantification of fluorescence intensities. **E.** Stills from live imaging of CB-EME expressing parasites co-stained with Mitotracker (mitochondria). The branched mitochondrial structure (DMSO) is not disrupted upon loss of PfACT1 (RAP). See also **Video V7**. **F.** IFA showing apicoplasts (red) colocalised with branches of the F-actin network (CB-EME, DMSO) and the disruption of the network together with apicoplasts when PfACT1 is deleted (RAP). Scale bar 5μm.

We reasoned that since most canonical actin filament stabilising and nucleating proteins are absent in Apicomplexa, the parasite must depend on formins for F-actin assembly. Previously, PfFRM1 has been localised to the invasion junction and PfFRM2 to the cytosol (Baum, Tonkin et al. 2008). Since we observed the intracellular F-actin network in the cytosol, we speculated that Formin-2 is the main nucleator of F-actin during intracellular parasite development, even though it has been implicated in host cell invasion in the case of *T.gondii* (Daher, Plattner et al. 2010).

### 2. Apicomplexan Formin-2 sequences contain a PTEN-C2-like domain found usually in plant formins

Formins possess a formin homology (FH) 1 and an FH2 domain, which nucleate actin monomers as well as elongate unbranched F-actin by continuous processive binding to the barbed end of the filament (Courtemanche 2018). In a previous report (Baum, Tonkin et al. 2008), only FH1/FH2 domains were described for apicomplexan formins. Here, we queried for presence of known PFAM domains using NCBI conserved domain search and in addition to FH1/FH2, found tetratricopeptide repeat (TPR) domains in both PfFRM1 and TgFRM1, while a PTEN C2-like domain was recognised in PfFRM2 and TgFRM2 (**Fig. 4A**). This led us to hypothesise that Formin-1 and Formin-2 with different N-terminal domains diverged early in evolution and different domain organisations have been retained for different functions. We queried for various FH2-domain containing proteins from Apicomplexans and found that Formin-2-like sequences are found in a different clade from Formin-1-like sequences (**Fig. 4B**), as also previously noted (Baum, Tonkin et al. 2008). Strikingly, the PTEN-C2-domain (or a diverged PTEN-C2 domain) was found only in Formin-2-like sequences (**Fig. 4B**). Interestingly, PTEN-C2 domains are important for membrane recruitment (Das, Dixon et al. 2003) and it has been shown to be recruited to rice chloroplast membranes (Zhang, Zhang et al. 2011), leading us to hypothesise that a similar mechanism operates for apicoplast recruitment of Formin-2 sequences in apicomplexans.

**Figure 4.**
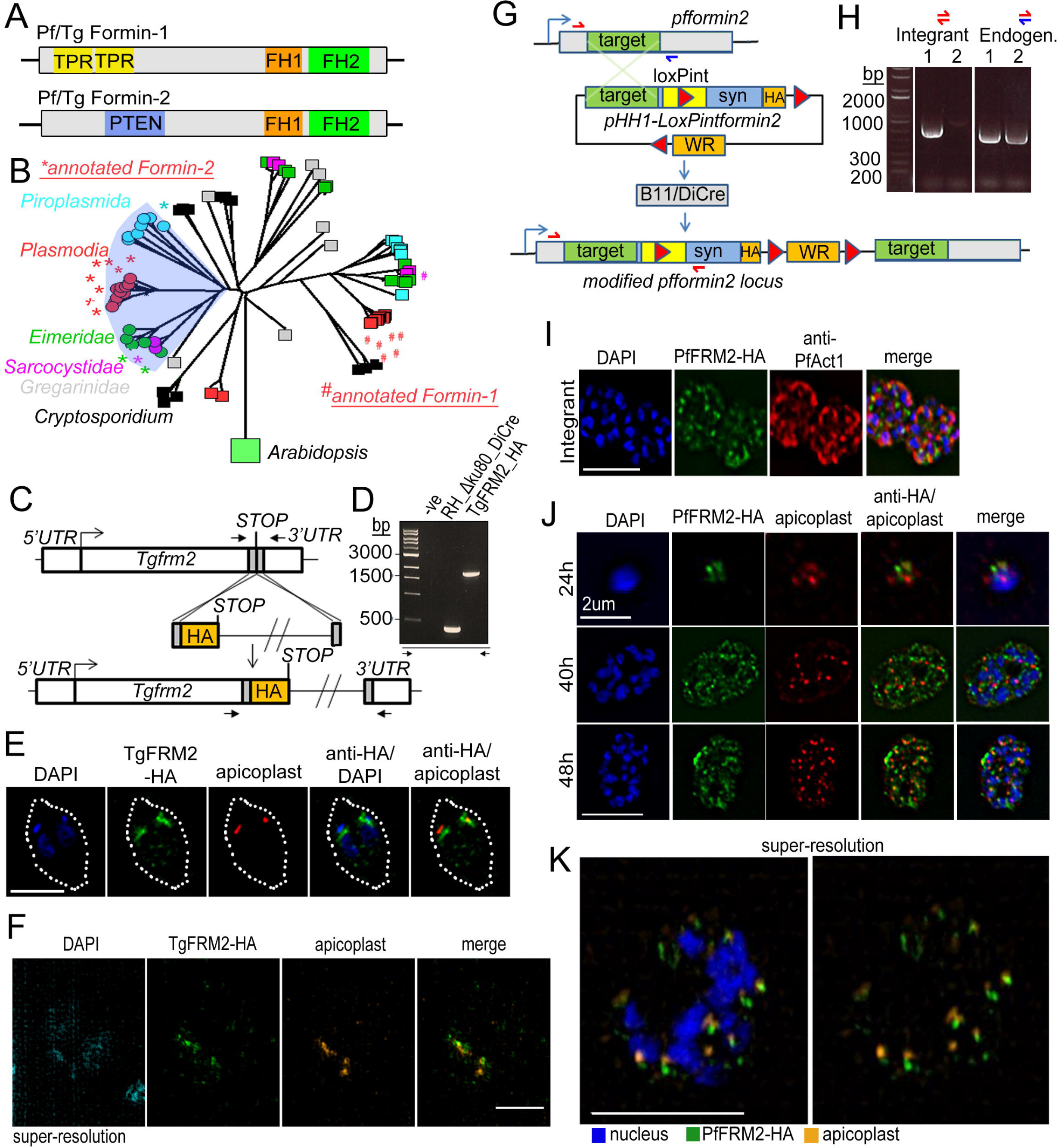
Apicomplexan formins have distinct protein domains, with Formin-2 localising to apicoplasts in *Toxoplasma* and *P. falciparum*. **A.** Other than the conserved FH1/FH2 domains, Pf/Tg Formin-1 contain tetratricopeptide repeat domains, while Pf/Tg Formin-2 contain a PTEN C2-like domain. **B.** Rooted neighbour-joining tree of FH2 domains detected in apicomplexan sequences flagged by hmmsearch and extracted from alignments produced by hmmalign, both using the PFAM profile PF02181.23. Proteins with sub-sequences similar to PTEN-C2 domains (detected by psi-Blast) are indicated with circular leaf symbols (and shaded blue). Those sequences annotated as Formin-1 (#) and Formin-2 (*) are indicated. Colour coding of the leaf nodes: Red: Plasmodium, Green: Eimeria, Magenta: Sarcocystidae, Cyan: Piroplasmida, Black: Cryptosporidium, Grey: Gregarinidae **C.** Strategy depicting endogenous C-terminal HA tagging of TgFRM2 in Toxoplasma. CRISPR/Cas9 was exploited to introduce a double-stranded DNA break and repair DNA amplified by PCR with homologous DNA regions coding for 3xHA. **D.** Diagnostic PCR confirming integration of DNA described in C into the RH_Δku80_DiCre line. **E.** IFA showing localisation of TgFRM2-HA (green) at the vicinity of the apicoplast staining (anti-G2Trx, red). Nuclei are stained with DAPI (blue). White dotted line depicts the parasite vacuole outline. Toxoplasma parasites were fixed 24h after inoculation. **F.** Super-resolution microscopy confirming the close apposition of TgFRM2-HA (green) to the apicoplast (anti-G2Trx, orange). Toxoplasma parasites were fixed 24h after inoculation. Scale bar is 2.5μm. **G.** Strategy showing simultaneous floxing and C-terminal HA tagging of the *pfformin2* locus using single cross over recombination into a DiCre expressing strain to give rise to the LoxPpfformin2 strain (modified). Primers for diagnostic PCR have been annotated as half arrows. **H.** Diagnostic PCR confirming integration in one of the two transfected lines (integrant). **I.** IFA showing localisation of PfFRM2-HA (green) in the context of a PfACT1-antibody staining (red). Nuclei are stained with DAPI (blue). **J.** IFA showing localisation of PfFRM2-HA adjacent to the apicoplast using a CPN60 antibody (red) throughout *P. falciparum* intracellular development (20, 40, 48h). K. Superresolution image confirming the tight apposition of PfFRM2 (green) with apicoplasts (orange). Scale bars are 5μm, except where stated otherwise.

### 2. *Plasmodium* and *Toxoplasma* Formin-2 localise adjacent to apicoplasts

In order to characterise the role of Formin-2 in detail, we decided to perform a comparative analysis in both *T.gondii* and *P. falciparum*. Therefore, we epitope tagged Formin-2 in both parasites. For tagging in *T.gondii* we used a CRISPR/Cas9-based strategy to introduce a 3x hemagglutinin (3HA) tag at the TgFRM2 C-terminus (**Fig. 4C**) and confirmed correct integration by diagnostic PCR (**Fig 4D**). Upon colocalisation with the anti-apicoplast antibody G2-Trx (Biddau and Sheiner, unpublished), we found TgFRM2 to be localised adjacent to apicoplasts (**Fig, 4E**), which was confirmed by superresolution microscopy (**Fig. 4F**). For localisation of PfFRM2, we simultaneously epitope tagged and floxed PfFRM2 by single cross-over homologous recombination in a DiCre-expressing parasite strain (**Fig. 4G**) and confirmed integrants by diagnostic PCR (**Fig. 4H**). Integrants were cloned by limiting dilution and two distinct clones of ‘LoxPpfformin2’ were used for phenotypic characterisation. PfFRM2 showed a punctate pattern in context of PfACT1 antibody staining (**Fig 4I**). Next, we checked for PfFRM2 localisation in relation to the apicoplast and observed a close apposition of the apicoplasts with the PfFRM2 staining throughout the 48h malaria life-cycle (**Fig. 4J**), which was confirmed by super-resolution microscopy (**Fig. 4K**). In conclusion, both *Toxoplasma* and *P. falciparum* Formin-2 localises adjacent to apicoplasts.

### 3. DiCre-mediated conditional disruption of Formin-2 causes a defect in apicoplast segregation in *P. falciparum*

Next, we wished to evaluate the fate of *P. falciparum* upon conditional DiCre-mediated disruption of the *pffrm2* gene (**Fig 5A**). 1h old tightly synchronised ring stage parasites were divided into two flasks and either pulse-treated with rapamycin (RAP) or DMSO (control) for 4h and their phenotype determined at 44h post RAP-treatment. Excision was determined by diagnostic PCR of the genomic locus (**Fig 5B**) and fitness of the PfFRM2 conditional knock out (KO) was measured by a growth curve which showed significant loss of viability (**Fig. 5C**). Loss of protein was ~90% as determined by Western blot (**Fig. 5D**) and was confirmed by IFA (**Fig. 5E**), which indicated a loss of protein in ~95% parasites (N=350). Giemsa stained PfFRM2 KO parasites were dysmorphic with apparent inclusions of haemoglobin (**Fig. 5F**, **red arrows**). In order to determine the morphological defects in FRM2 KO parasites, we co-stained PfFRM2 KO parasites with several organellar markers and were unable to see significant differences (not shown), except for apicoplast segregation (**Fig. 5G**). The number of parasites with normally segregated apicoplasts was significantly reduced, with a high percentage of cells showing collapsed or morphologically aberrant apicoplasts (**Fig 5G, H**). A range of apicoplast phenotypes was evident, from totally collapsed, intermediate to apparently normal (**Fig S3A**). To determine if the loss of viability of the PfFRM2 KO parasites was solely due to loss of the apicoplast, we attempted to rescue the phenotype with 200μM isopentenyl pyrophosphate (IPP) which has been previously shown to complement growth in parasites lacking apicoplasts (Yeh and DeRisi 2011). However, we did not see any improvement in viability, indicating that the loss of fitness is due to additional defects caused by abrogation of F-actin dynamics in the parasite. We determined number of nuclei in 44h PfFRM2 KO parasites and found a significant decrease in the number of nuclei (**Fig. S3B**), indicating a developmental defect. Since PfACT1 is required for normal cytokinesis (Das, Lemgruber et al. 2017), we interrogated if the IMC is normally formed in PfRM2 KO parasites. We purified mature schizonts on a 70% Percoll cushion and determined by IFA that IMC formation was compromised in these parasites, with normal IMC staining dropping from 58±8% in WT to 19±8% in PfFRM2 KOs (**Fig. S3C, D**). When we allowed PfFRM2 KO parasites to egress and compared them to control parasites, we found conjoined merozoites in PfFRM2 KOs, a defect previously seen in PfACT1 KO parasites (Das, Lemgruber et al. 2017), indicating that PfFRM2 and PfACT1 coordinate cytokinesis in *P. falciparum*.

**Figure 5.**
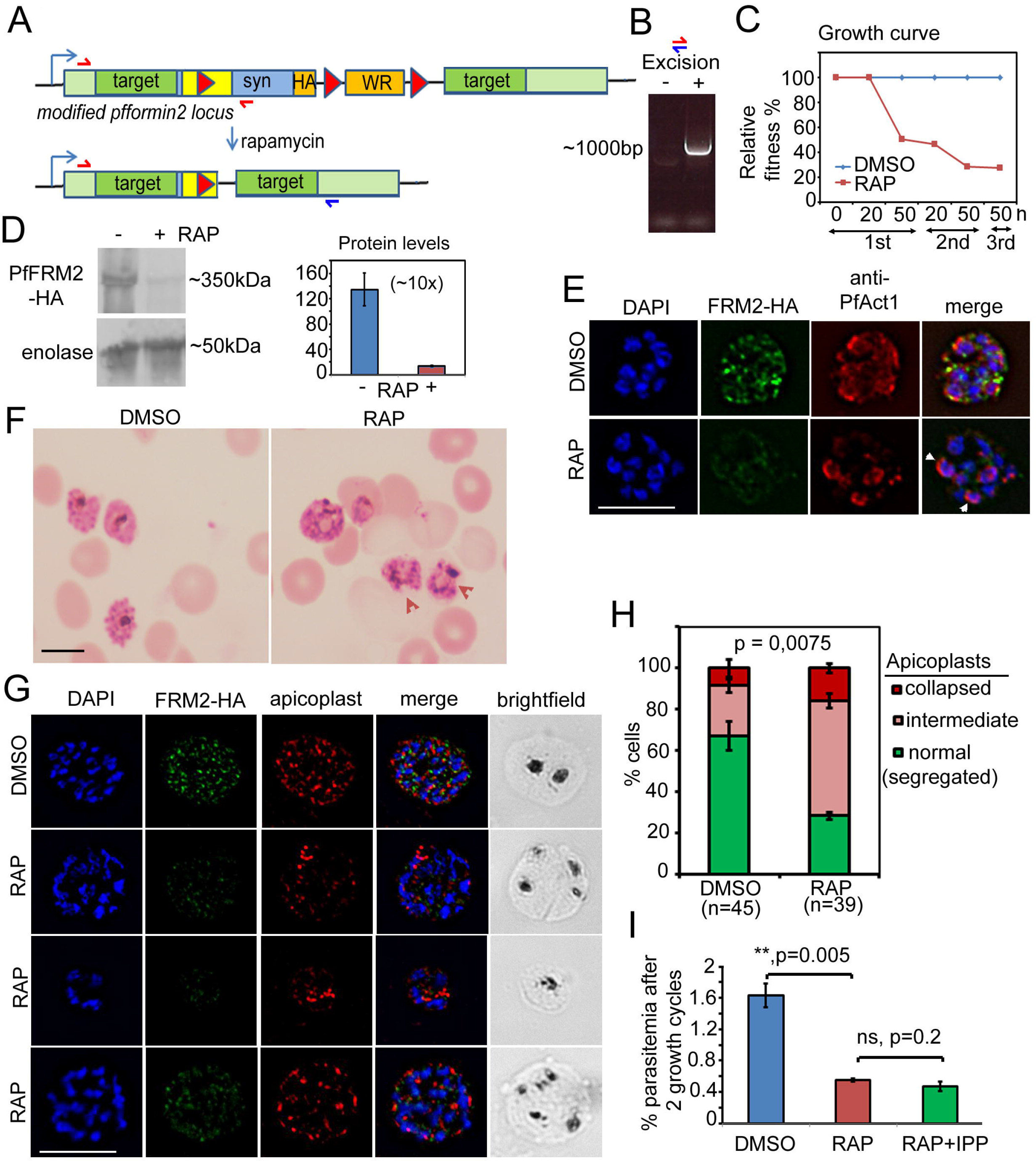
Conditional deletion of PfFRM2 disrupts apicoplast segregation and causes a severe fitness defect. **A.** Strategy showing the DiCre-mediated genomic excision of the LoxPpfFRM2 locus. Primers for diagnostic PCR have been annotated as red/blue half arrows. **B.** Diagnostic PCR confirming genomic excision of the *pffrm2* locus upon rapamycin treatment (+). **C.** A growth curve showing the relative fitness of RAP-treated PfFRM2 KO parasites in comparison to DMSO controls. Various time points from the pulse treatment of 1h-old rings at time 0 in the 1^st^, 2^nd^ and 3^rd^ growth cycles have been measured. **D.** left panel, Western blot showing the loss of PfFRM2-HA upon RAP-treatment, enolase has been used as a control. Right panel, Quantification of PfFRM2-HA from three different blots shows at least a 10-fold drop in protein levels, Error bars depict SD. **E.** IFA showing loss of PfFRM2-HA staining (green) upon RAP-treatment. Levels of PfACT1-staining (red) do not change. **F.** Giemsa stained images of RAP-treated parasites reveal dysmorphic parasites. **G.** Apicoplast segregation (red) is affected to various degrees in RAP-treated parasites as compared to DMSO controls. See also **Figs. S3A** and **6D**. **H.** Quantification of phenotypes seen in **G.** Error bars depict SD. I. Isopentenyl pyrophosphate (IPP) cannot rescue the fitness defect (RAP+IPP) in PfFRM2 KO parasites (RAP) as compared to parasitemia to the DMSO controls. Error bars depict SD. Scale bars 5 μm.

### 4. DiCre-mediated disruption of Formin-2 abrogates the actin network in *P. falciparum* trophozoites

Next, we wished to determine whether F-actin assembly and dynamics is interfered upon deletion of PfFRM2 and if this directly affects apicoplast segregation. We expressed CB-EME in LoxPpfformin2 to generate the line LoxPpfformin2/CBEME (**Fig. 6A**) and visualised actin filaments (**Fig 6B, DMSO**). As observed in wild-type parasites (**Fig 1**), F-actin was decorated with punctate PfFRM2-HA staining (Fig S4, **Pf**). Upon DiCre-mediated excision in ring stages, we saw a complete abrogation of the dynamic F-actin network in mature schizont stage parasites (**Fig 6B RAP**), which dropped from exhibiting an F-actin network in 92±5% cells in WT to 5±4% cells in PfFRM2 KO (**Fig. 6C**). Furthermore, we confirmed the apicoplast phenotype in parasites expressing CB-EME (**Fig 6D**).

**Figure 6.**
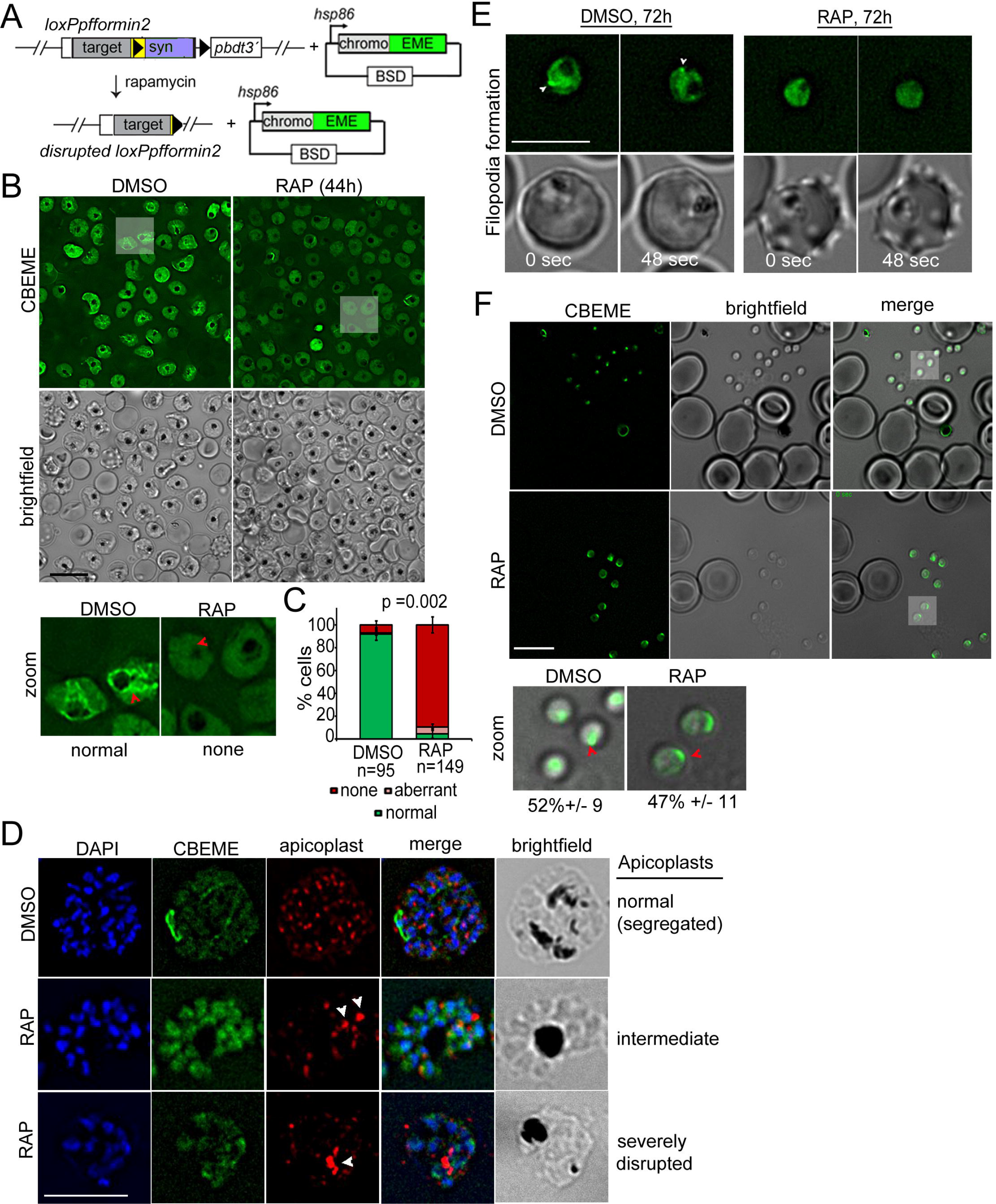
Conditional deletion of PfFRM2 abrogates the intracellular F-actin network but not the apical polymerisation of F-actin prior to invasion. **A.** Strategy showing episomal expression of CB-EME in the LoxPpffrm2 background. **B.** Upper panels: Stills from a video showing loss of normal intracellular F-actin fluorescence (green). Brightfield images have been provided. See also Video V8. Lower left panel: Zoomed images of indicated parasites in upper panels showing loss of the actin network in RAP (none, red arrows) as compared to DMSO controls (normal). **C.** Graph showing loss of normal F-actin fluorescence in ~95% RAP-treated parasites. >90% of DMSO controls show presence of the network. **D.** IFA staining of the apicoplast with a CPN60 antibody (red) on top of the fluorescent F-actin network (green) shows a defect in apicoplast segregation in RAP treated parasites (white arrows). Nuclei are stained in blue and brightfield images are provided alongside. Examples of normally segregated apicoplasts, intermediate and severely disrupted apicoplasts have been provided. **E.** Loss of filopodia in RAP-treated young trophozoites imaged 72h post-RAP treatment. Stills from a video have been shown, where time points are depicted in white. Normal filopodia are marked with white arrows in DMSO controls. **F.** Post-schizont egress, merozoites from DMSO controls and RAP-treated group show similar propensity to polymerise apical F-actin (CB-EME fluorescence shown in green). Red arrows show apical F-actin in zoomed images (lower panels). Scale bars 5μm.

Interestingly, we observed that actin-rich filopodia formed at the cell-periphery (see also **Fig. 2A, B**) disappeared upon PfFRM2 disruption (**Fig. 6E**), providing a mechanism for this novel phenomenon.

Since PfFRM1 was localised to the parasite apex/ invasion junction during host cell entry (Baum, Tonkin et al. 2008), we reasoned that apical polymerisation of F-actin should not be affected in PfFRM2 KO parasites, if indeed the two formins perform distinct functions in their distinct localisations. To this end, we allowed mature schizonts to egress and release free merozoites and subsequently imaged them by live fluorescence microscopy. Consistent with this hypothesis, we found that the ability of F-actin polymerisation at the parasite apex was not compromised in PfFRM2 KO parasites (**Fig. 6F**), strongly indicating distinct functions of PfFRM1 and PfFRM2.

### 5. Conditional deletion of Formin-2 in*Toxoplasma* disrupts apicoplast segregation and F-actin dynamics

Finally, in order to assess if the function of Formin-2 is conserved in apicomplexan parasites, we analysed its role in *T.gondii*. We first checked the localisation of TgFRM2 with respect to the F-actin network. Similar to *P. falciparum*, TgFRM2-HA formed puncta on CB-EME labelled F-actin network within the parasite, where it appeared to co-localise with a polymerisation centre (**Fig. S4 Tg**). Next, we simultaneously floxed *tgfrm2* together with addition of C-terminal YFP tag to create the LoxPTgFRM2 line (**Fig. 7A**). This enabled us to confirm localisation of Formin-2 and determine the comparative effect of a conditional KO in *Toxoplasma*. Integration of the C-terminal YFP-tag and *LoxP* sites was confirmed by diagnostic PCR, as was excision of the *tgfrm2* locus upon RAP-treatment (**Fig S5A**). For the localisation of TgFRM2 it was necessary to stain fixed parasites with a YFP-antibody, suggesting low expression levels of TgFRM2. We confirmed localisation of TgFRM2 adjacent to the apicoplast (**Fig 7B upper panel**). Upon RAP-treatment, excision of TgFRM2 was apparent in 14% (clone A) or 33% (clone B) of parasites, as assessed by quantification of parasites where no TgFRM2 could be detected by IFA. Importantly, loss of TgFRM2 staining correlated with an apicoplast segregation phenotype in the majority of parasites (**Fig 7B lower panel**, **42.4 % [n=66] in clone A and 70% [n=100] in clone B**). A baseline apicoplast segregation phenotype was observed in 2% (clone A) or 1% (clone B) of vacuoles in the control population. Transient expression of CB-EME in LoxPTgFRM2 parasites enabled us to image F-actin and demonstrated that, in good agreement with data from *P. falciparum*, intracellular F-actin was adjacent to the apicoplast (**Fig 7C control**). Intriguingly, excision of TgFRM2 (**Fig. 7C RAP**) led to the disappearance of intracellular F-actin, while (in contrast to *P. falciparum*), the intravacuolar F-actin network was still present (Fig.7C), indicating that another formin, potentially Formin-3 (Daher, Klages et al. 2012), which is not present in *P.falciparum*, contributes to its formation.

**Figure 7.**
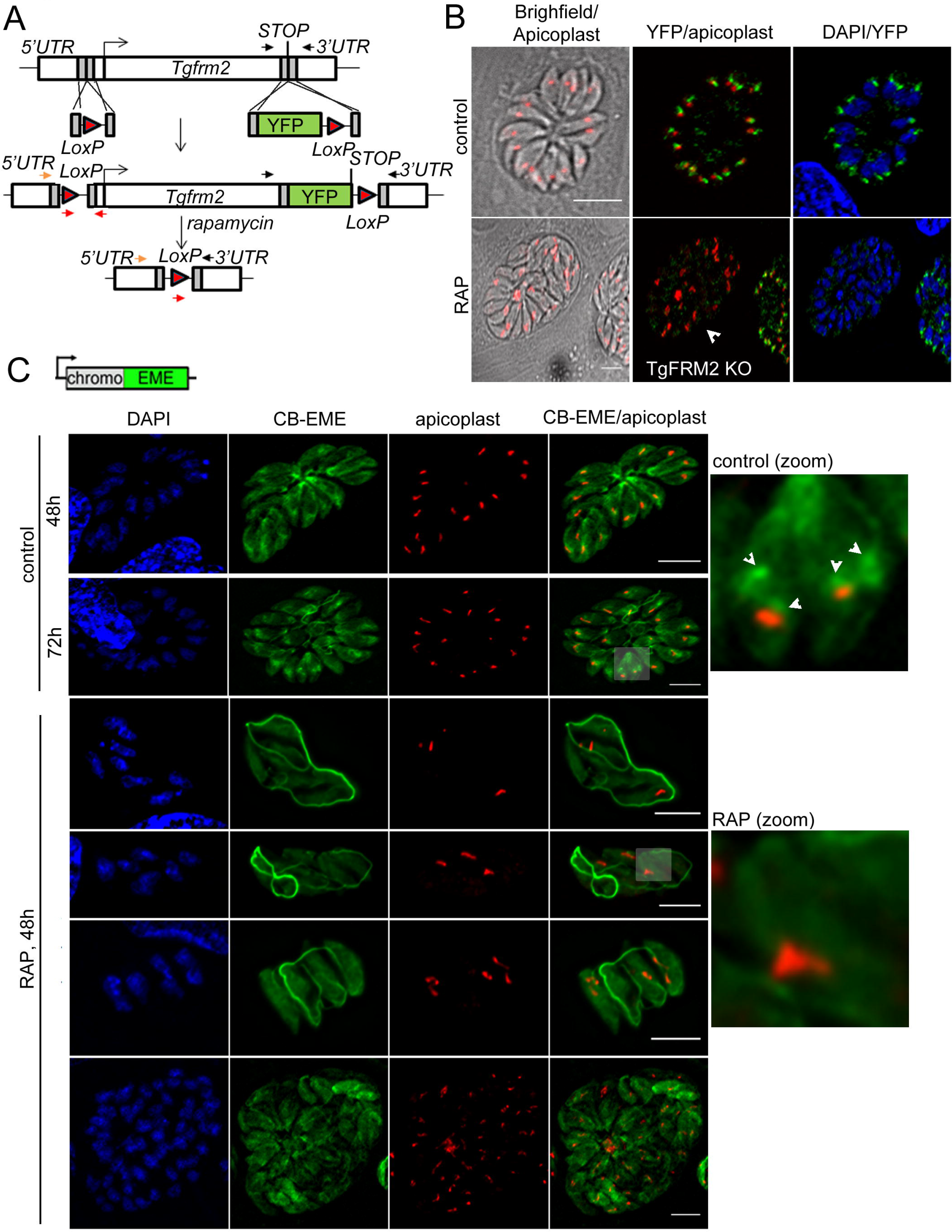
Conditional deletion of TgFRM2 disrupts normal segregation of apicoplasts together with abrogation of the intracellular F-actin polymerisation centre. **A.** Strategy to generate LoxPTgFRM2-YFP, a floxed and C-terminal YFP-tagged *TgFRM2* locus in the RH_Aku80_DiCre line. For this purpose, CRISPR/Cas9 was exploited to introduce DNA double-strand breaks in the 5’ UTR and C-terminus of the *TgFRM2* gene. Integration was confirmed by PCR (see **Fig. S5A**). Arrows represent PCR primers used in **Fig. S5A**. **B.** IFA staining with anti-YFP (TgFRM2-YFP) and anti-Atrx1 (apicoplast) shows an apicoplast segregation defect in TgFRM2-YFP KO parasites. In control parasites, TgFRM2-YFP localises to the vicinity of the apicoplast (upper panel). The loss of TgFRM2-YFP causes an apicoplast segregation defect (lower panel, white arrow). The lower panel depicts a TgFRM2-YFP KO vacuole together with a TgFMR2-YFP positive vacuole for comparison. Scale bars are 5μm. **C.** IFA depicting CB-EME and apicoplast (anti-CPN60) in control and RAP-treated LoxPTgFRM2-YFP parasites. In untreated parasites, the apicoplast localises to intracellular actin polymerisation centres (control, white arrows in zoom). Parasites exhibiting TgFRM2 KO-specific apicoplast phenotype lack intracellular actin polymerisation centres. Zoomed images depict indicted areas. Scale bars are 5μm. IFAs depicted in this figure were performed for the LoxPTgFRM2 clone B.

## Discussion

Due to the unconventional behaviour of apicomplexan actin, the visualisation of actin filaments in *P. falciparum* was hampered by the lack of reagents and F-actin sensors that do not interfere significantly with F-actin polymerisation and depolymerisation. Therefore, previous attempts to use established indicators such as Life-Act from other eukaryotic systems failed (Tardieux 2017). In a recent study it was shown that actin-binding nanobodies fused to epitope tags could be expressed in *Toxoplasma gondii*, allowing for the first time to analyse F-actin localisation and dynamics in living parasites (Periz, Whitelaw et al. 2017). Here we successfully adapted this technology to *P. falciparum* using two different epitope tags, the halo tag and the emerald tag. This allowed us to visualise F-actin throughout the asexual life-cycle of *P. falciparum* and in gametocytes without causing any aberrant phenotypes, suggesting that this reagent, as in the case of *T.gondii* (Periz, Whitelaw et al. 2017) and all other eukaryotes tested so far (Melak, Plessner et al. 2017) does not significantly interfere with F-actin dynamics. Importantly, validation of this reagent using either F-actin modulating drugs or a conditional mutant for PfACT1 led to expected results and phenotypes, demonstrating that F-actin dynamics is finely balanced in the parasite. Since expression of chromobodies does not interfere with parasite viability, it can be assumed that its influence on F-actin dynamics is at best minor, potentially leading to slightly altered F-actin dynamics. Similar influences of F-actin sensors have been observed and discussed in other eukaryotes (Melak, Plessner et al. 2017). As a general rule, any F-actin sensor will influence F-actin dynamics or might not be able to stain all F-actin structures. In this respect, F-actin binding nanobodies are a novel tool and appear to have only minimal effects on F-actin dynamics compared to other sensors (Melak, Plessner et al. 2017), which might explain that they are well tolerated in Apicomplexans, where rapid F-actin dynamics appears to be critical for parasite viability. Using this novel tool, we not only demonstrate here that F-actin can be found in very close proximity of apicoplasts, we also demonstrate a novel function of F-actin during the intracellular development of *P. falciparum*: formation of highly dynamic, filopodia-like structures. While the role of this process is currently unknown, we show that interference with F-actin dynamics, either by addition of actin-modulating drugs of conditional mutants for PfACT1 or PfFRM2, led to a complete blockade in the formation of these structures.

Super-resolution imaging revealed a complex F-actin network in *P. falciparum*, similar to that observed in *T.gondii* (Periz, Whitelaw et al. 2017) with extensive filaments around the FV. Importantly, our previous characterisation of a conditional mutant for PfACT1 highlighted the diverse functions of actin during the asexual life cycle of the parasite (Das, Lemgruber et al. 2017), which perfectly correlates to the localisation and dynamics found here using chromobody-expressing parasites. PfACT1 is essential for *P. falciparum* invasion into erythrocytes and we show for the first time the temporal and spatial dynamics of actin polymerisation by live microscopy prior to invasion. Despite a growing body of evidence suggesting the importance of calcium signalling and phosphorylation of IMC proteins by kinases such as CDPK1 and PKA (Baker, Drought et al. 2017, Kumar, Kumar et al. 2017) during invasion, what triggers the polymerisation of actin is largely unknown. Our data suggest that early signalling events just after egress are a trigger for actin-polymerisation at the apical end. This is likely to be mediated by an apically resident nucleator of F-actin, a likely candidate being Formin-1, since PfFRM2 KO parasites could still polymerise actin at the apical end, as demonstrated in this study.

Therefore, the expression of chromobodies in *P. falciparum* allows us to phenotypically probe the state of the F-actin network *in vivo* in a rapid and robust manner. F-actin can be clearly visualised during growth, in invading merozoites and in gametocytes - opening up many avenues for further research. Using this novel tool, combined with powerful reverse genetics made possible by the DiCre system (Collins, Das et al. 2013) we show here that Formin-2 in both *Toxoplasma* and *Plasmodium* is required for the intracellular polymerisation of F-actin, a mechanism employed by the parasite for correct segregation of apicoplasts. Using extensive bioinformatic searches within alveolates, we found the presence of a PTEN-C2-like domain only in apicomplexan Formin-2 sequences. This domain has been demonstrated in rice to be responsible for Formin-2 targeting to chloroplast membranes (Zhang, Zhang et al. 2011). It is therefore likely that the apicomplexan PTEN-C2-like domain is used for apicoplast recruitment of apicomplexan Formin-2.

Interestingly, while we found that the role of Formin-2 appears to be conserved in *T.gondii* and *P. falciparum* for intracellular F-actin dynamics and apicoplast inheritance (**Fig. 8**), in the case of *T. gondii* the intravacuolar F-actin network is still formed, suggesting that *T.gondii* and potentially other coccidia have additional, compensatory mechanisms at their disposal to form this network, which appears to be critical for material exchange, synchronised replication of parasites and host cell egress (Periz, Whitelaw et al. 2017).

**Figure 8.**
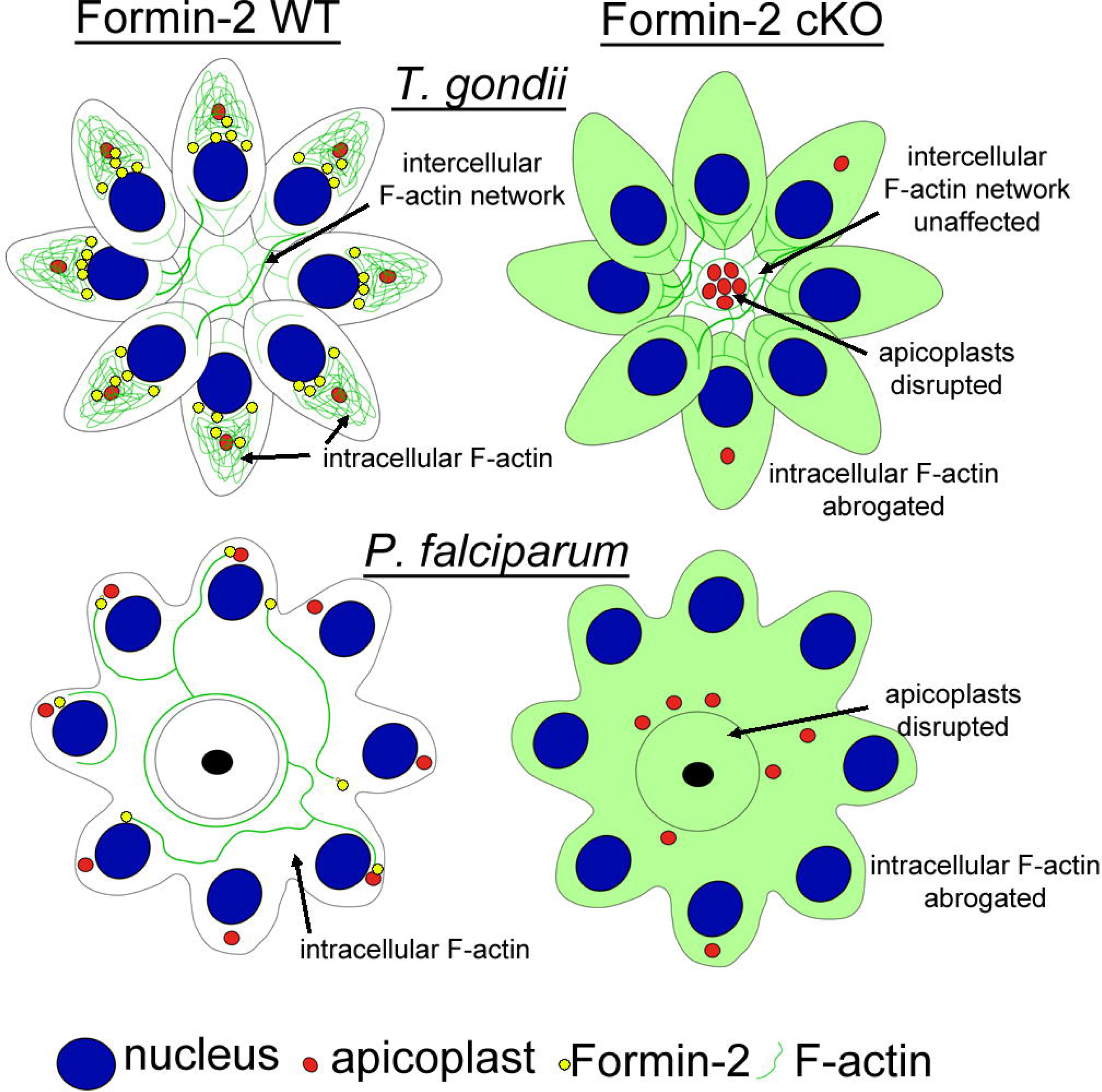
A model showing the dependence of apicoplast segregation on the intracellular F-actin network in *P. falciparum* and *Toxoplasma*. In apicomplexan parasites, despite using different replication modes (endodyogeny, schizogony), the function of actin and Formin-2 is highly conserved with regards to intracellular F-actin dynamics and apicoplast inheritance. One difference appears to be the formation of an intravacuolar network that can still occur in the case of *T.gondii* but not *P. falciparum* upon disruption of Formin-2.

In conclusion, we show here that chromobodies can be used to determine F-actin dynamics in apicomplexan parasites and will form the basis for functional *in vivo* studies of other actin regulatory proteins found in apicomplexans. Furthermore, they can be used as an efficient tool to probe for drugs specifically interfering with F-actin dynamics in apicomplexans.

## Experimental procedures

### 1. Culture and transfection of *P. falciparum*

*P. falciparum* was cultured in RPMI 1640 with Albumax (Invitrogen) and schizonts were purified on a bed of 70% Percoll as described previously (Blackman 1994). About 10 μg of plasmid was ethanol precipitated and resuspended in 10 μL sterile buffer TE (Qiagen). The Amaxa™ P3 primary cell 4D Nucleofector™ X Kit L (Lonza) was used for transfections. The input DNA was added to 100 μL P3 primary cell solution, mixed with 10-20 μL of packed synchronous mature schizonts and added to the cuvette, which was electroporated in a 4D-Nucleofector machine (Lonza) using the program FP158. The transfected schizonts were rapidly added to 2 mL of complete medium (RPMI with Albumax supplemented with glutamine) containing erythrocytes at a haematocrit of 15%, and left shaking in a shaking incubator at 37°C for 30 min. Finally the cultures were supplemented with 7 mL of complete RPMI medium to obtain a final haematocrit of 3% and incubated overnight at 37°C in a small angle-necked flask (Nunc™). Parasites were selected by use of appropriate drug medium. The culture medium was subsequently exchanged every day for the next 4 days to remove cell debris which accumulates during electroporation and then twice a week until parasites were detected by Giemsa smear. Drug-resistant parasites were generally detectable in thin blood films 2-3 weeks post-transfection. After this, parasite stocks (at ~5% ring parasitaemia) were cryopreserved in liquid nitrogen. Lines were then cloned by limiting dilution using a simple plaque assay (Thomas, Collins et al. 2016).

### 2. Cloning and expression of chromobodies in *P. falciparum*

The CB-HALO and CB-EME plasmid consists of a sequence encoding actin chromobody from Chromotek followed downstream by an in frame sequence encoding Halo (Promega) or the emerald tag. CB-EME and CB-HALO sequences were amplified by PCR and cloned into the vector pB-map2gfpdd (Nicholas Brancucci, unpublished) via restriction sites NheI and HindIII to remove the map2gfpdd sequence and put the CB-sequences under the *hsp86* promoter. The resulting plasmids pB-CBEME and pB-CBHALO were sequenced on both strands to confirm correct nucleotide sequences. These constructs were transfected as described into the loxPACT1 parasite clone B2 (Das, Lemgruber et al. 2017) to obtain parasite lines LoxPPfACT1/CBEME, LoxPPfACT1/CBHALO and into the parental 1G5DiCre clone (Collins, Das et al. 2013) to obtain the line CBEME/1G5DiCre and CBHALO/1G5DiCre. Lines were selected with 2.5 μg/mL blasticidin. CB-EME expression was visible by fluorophore excitation/emission in the green range and the HALO ligand was made visible by use of the ligand HALO-TMR at 1:40,000 with excitation/emission in the red range. Alternatively antibodies were used against the HALO tag to stain for CB-HALO.

### 3. *P. falciparum* IFA

Thin blood films were made on glass slides and fixed in 4% paraformaldehyde in PBS for 20 min. The slides were then permeabilised with 0.1% Triton-X/PBS for 10 min, washed and blocked overnight in 4% BSA/PBS. Antigens were labelled with suitable primary and secondary antibodies in 4% BSA/PBS with 5 min PBS washes in between. Slides were finally air dried and mounted with DAPI-Fluormount-G^®^ (SouthernBiotech).

Staining of the RON4 junction in CB-EME expressing was performed by fixation and immunostaining in solution as described previously (Riglar, Richard et al. 2011).

For image acquisition, z-stacks were collected using a UPLSAPO 100x oil (1.40NA) objective on a Deltavision Core microscope (Image Solutions - Applied Precision, GE) attached to a CoolSNAP HQ2 CCD camera. Deconvolution was performed using SoftWoRx Suite 2.0 (Applied Precision, GE).

An Elyra S1 microscope with Superresolution Structured Illumination (SR-SIM) (Zeiss) was used for super-resolution imaging.

### 4. Time lapse microscopy of live *P. falciparum*

Video microscopy of *P. falciparum* schizont egress was performed as described previously (Collins, Hackett et al. 2013). Synchronised schizonts were Percoll^®^ purified and treated with 1 μM C2 in RPMI medium with Albumax^®^ (Gibco) for 4h. Microscopy chambers (internal volume ~80 μl) for observing live schizonts were built by adhering 22×64 mm borosilicate glass coverslips to microscope slides with strips of double-sided tape, leaving ~4 mm gaps at each end. C1 was washed off before video microscopy and the schizonts were immediately resuspended into warm (37°C) RPMI (with Albumax) and introduced by capillary action into the pre-warmed chamber. The chamber was transferred to a temperature-controlled microscope stage at 37°C on a Deltavision Core microscope (Image Solutions - Applied Precision, GE). Images were routinely collected at 5 s intervals, beginning 6 min 30 sec after washing off C1, over a total of 30 min.

Other than during egress, CB-EME and CB-HALO expressing parasites were imaged at intervals of 1 sec.

### 5. Bioinformatics

Proteomes of interest (Table T2) were downloaded from the UniProt-KB website (www.uniprot.org). These were concatenated into a single proteome sequence dataset. All sequence identifiers and annotations referred to are from UniProt Hidden Markov Models (PFAM profiles) PF02181.23 (FH2.hmm, Formin Homology 2 Domain) and PF10409.9 (PTEN_C2.hmm, C2 domain of PTEN tumour-suppressor protein) were downloaded from Pfam (El-Gebali, Mistry et al. 2018). These profiles were used with the HMMER package (HMMER 3.1b1 (May 2013); http://hmmer.org/) to search the proteome sequences (hmmsearch), and to align sequences of interest (hmmalign). The proteome sequence dataset was searched for FH2 domains (FH2.hmm) with hmmsearch, and sequences with regions scoring >28bits recorded. These sequences were retrieved from the dataset, and subjected to alignment against the FH2.hmm. The profile conformant subsequences were extracted from the alignment and this sequence set subjected to alignment using: 1) hmmalign to FH2.hmm, 2) clustalw (Thompson, Higgins et al. 1994) 3) muscle (Edgar 2004) and 4) T_Coffee. These multiple sequence aligments were combined and evaluated in T_coffee (Keller, Kollmar et al. 2011) using the-aln and-special_mode evaluate options of T_coffee and the alignment edited to remove columns of avgerage quality <4 and occupancy <30% (T_coffee - other_pg seq_reformat option). Rooted neighbour-joining trees of Formin Homology type 2 domains (FH2) was contructed from this alignment (or subsets of it) using the SplitsTree program [version 1.14.8,*]. The proteome dataset was searched for the presence of PTEN_C2 conformant sequences. As only an inconsistent subset of sequences were found in both PTEN_C2 and FH2 selected sequences; one such subsequence (A0A1A7VGT3_PLAKH, residues 1096-1238) was used as the query of an iterative psi-blast [@], (E-value cutoff =10) using the proteome data set as the database. The program converged after 3 iterations. The sequences flagged by psi-blast as having PTEN_C2-like sequence were compared with the sequences flagged by hmmsearch as having FH2 domains, and such sequences annotated on the phylogenetic tree.

**Table T1.** Antibodies used in this study.

| Antibody | Reference |
| --- | --- |
| Anti actin | (Angrisano, Riglar et al. 2012), RRID: AB_2665920 |
| Anti CPN60 (apicoplast) | (Agrawal, van Dooren et al. 2009) |
| Anti RON4 | (Richard, MacRaild et al. 2010) |
| Anti GAP45 | (Jones, Kitson et al. 2006) |
| Anti MTIP | (Jones, Kitson et al. 2006) |
| Anti-enolase | (Dutta, Tewari et al. 2018) |
| Anti-HA | Roche (Cat #118743100) |
| Anti-YFP | Abcam (Cat #ab6556) |
| Anti-Atrx1 | (DeRocher, Coppens et al. 2008) |
| Anti-G2Trx | Biddau and Sheiner, unpublished. |

### 6. Creation of LoxPPfFRM2-HA and LoxPPfFRM2/CBEME strains

To obtain conditional truncation of the *pffrm2* gene we used silent *loxP* sites within a heterologous *P. falciparum* intron loxPint (Jones, Das et al. 2016). We ordered from Geneart^®^ a ~800 bp targeting sequence followed by the LoxPint module in the context AATTGTAG-LoxPint-ATAGCTTT followed by a recodonised version of rest of the 3’ region of the gene together with a C-terminal 3HA tag. This ordered synthetic fragment was cloned into the pHH1-loxPMSP1 plasmid (Das, Hertrich et al. 2015) via restriction sites AflII and ClaI, replacing the msp1 sequence with *pffrm2*, giving rise to the plasmid pHH1-LoxPintFormin2 (**Fig. 4G**). This was transfected into the DiCre expressing strain B11 (Perrin, Collins et al. 2018) and integrants selected by cycling on and off the drug WR99210 (Jacobus Pharmaceuticals, New Jersey, USA). The integrant line LoxPPfFRM2 was cloned by limiting dilution and two clones used for phenotypic characterisation. The strain LoxPPfFRM2/CBEME was created by transfecting the pB-CBEME plasmid into a LoxPPfFRM2-HA clone line and transfectants selected using the drug blasticidin (Sigma).

### 7. Conditional truncation of *pfactl* and *pffrm2*

Various floxed parasite strains were synchronised by Percoll and sorbitol as previously described (Collins, Hackett et al. 2013). Briefly, schizonts were purified on a bed of 66% Percoll and allowed to reinvade into fresh erythrocytes for 1-2h. The remainder of the schizonts were removed by Percoll and the freshly invaded rings were subjected to 5% sorbitol for 7 min at 37°C to lyse any remaining schizonts. The tightly synchronised rings were divided into two flasks and pulse-treated for 4h at 37°C with 100 nM rapamycin or with 1% DMSO as control. The rings were then washed and returned to culture. Phenotypic analysis was performed primarily 44h post RAP-treatment unless stated otherwise.

### 8. Culturing of *Toxoplasma* parasites and host cells

Human foreskin fibroblasts (HFFs) (RRID: CVCL_3285, ATCC) were grown on tissue culture-treated plastics and maintained in Dulbecco’s modified Eagle’s medium (DMEM) supplemented with 10% foetal bovine serum, 2 mM L-glutamine and 25 mg/mL gentamycin. Parasites were cultured on HFFs and maintained at 37° C and 5% CO2. Cultured cells and parasites were regularly screened against mycoplasma contamination using the LookOut Mycoplasma detection kit (Sigma) and cured with Mycoplasma Removal Agent (Bio-Rad) if necessary.

### 9. Microscopy for *Toxoplasma*

Widefield images were acquired in z-stacks of 2 μm increments and were collected using an Olympus UPLSAPO 100x oil (1.40NA) objective on a Delta Vision Core microscope (AppliedPrecision, GE) attached to a CoolSNAP HQ2 CCD camera. Deconvolution was performed using SoftWoRx Suite 2.0 (AppliedPrecision, GE). Further image processing was performed using ImageJ software.

Super-resolution microscopy (SR-SIM) was carried out using an ELYRA PS.1 microscope (Zeiss) as described previously (Periz et al., 2017). Images were acquired using a Plan Apochromat 63x, 1.4 NA oil immersion lens, recorded with a CoolSNAP HQ camera (Photometrics), and analysed using ZEN Black software (Zeiss) and ImageJ software.

### 10. *Toxoplasma* IFA

For immunofluorescence analysis, HFF monolayers infected with *Toxoplasma* parasites were grown on coverslips and fixed at the indicated time points in 4% paraformaldehyde for 20 min at RT. Afterwards coverslips were permeabilised in 0.2% Triton X-100 in 1x PBS for 20 min, followed by blocking (3% BSA & 0.2% Triton X-100 in 1x PBS) for at least 30 min. The staining was performed using indicated combinations of primary antibodies for 1 h and followed by secondary Alexa Fluor 488 or Alexa Fluor 594 conjugated antibodies (1:3000, Invitrogen-Molecular Probes) for another 45 min, respectively. The primary antibodies used in this studies are anti-G2Trx (1:500, rabbit, apicoplast, Dr Lilach Sheiner), anti-HSP60/CPN60 (1:2000, rabbit, apicoplast, Dr Lilach Sheiner), anti-Atrx1 (1:500, mouse, apicoplast, Prof Peter Bradley), anti-HA (1:500, rat, Roche [cat# 1187431001]) and anti-YFP (1:500, rabbit, Abcam [ab6556

### 11. Generation of the TgFRM2-HA and loxPTgFRM2-YFP strains in RHΔku80DiCre parasites

Guide RNAs targeting the upstream region of TgFRM2 and the C-terminal region were designed using EuPaGDT (Ref: Duo Peng and Rick Tarleton. EuPaGDT: a web tool tailored to design CRISPR guide RNAs for eukaryotic pathogens. Microbial Genomics. 2015. doi: 10.1099/mgen.0.000033). These were cloned into a vector expressing a Cas9-YFP fusion as well as the specific gRNAs as previously described (Curt-Varesano, Braun et al. 2016). The designed gRNAs ACTTTTCATAGTATAGGAGG CGG and AATAGGGGTCTGTAGGTTAA GGG bind 989 bp upstream of the start codon and 12 bp upstream of the stop codon of TgFRM2 respectively. To introduce the upstream LoxP site, the LoxP sequence ATAACTTCGTATAGCATACATTATACGAAGTTAT flanked with respective 33 bp homology on each side was ordered as a 100 bp primer (ThermoFischer Scientific). The repair template for the C-terminal tag (HA or YFP) was generated by PCR using Q5 polymerase (New England Biolabs) from template plasmids with 50 bp of target-specific homology introduced via the primer. All tags are flanked by the same sequence, the upstream linker sequence GCTAAAATTGGAAGTGGAGGA encoding for the amino acid sequence AKIGSGG, the tag itself, a stop codon and the LoxP sequence. The YFP tag is superfolder YFP 2, and was sub-cloned from pSYFP2-C1 (gift from Dorus Gadella (Addgene plasmid # 22878; http://n2t.net/addgene:22878;RRID:Addgene_22878) (Kremers, Goedhart et al. 2006). All C-terminal repair templates were pooled, purified using a PCR purification Kit (Blirt). Together with 10 μg Cas9 vector encoding the respective gRNA, 1 × 107 of freshly released RHΔku80DiCre tachyzoites (an improved version created by Dr Moritz Treeck from the original (Andenmatten, Egarter et al. 2013)) were transfected using 4D AMAXA electroporation. 24 hours after transfection, parasites were mechanically released, filtered and sorted for transient YFP expression into 96 well plates using a FACS sorter (FACSARIA III, BD Biosciences). Individual plaques were screened by PCR and the C-terminus of TgFRM2 was sequenced (Eurofins Genomics). Into a clone with TgFRM2-YFP-LoxP, the upstream LoxP was introduced as described. Screening for upstream LoxP integration was performed by PCR with a primer binding at the junction of gRNA binding sequence and LoxP site. Using a different set of primers, the complete upstream LoxP site was amplified via PCR and verified by sequencing. Two distinct clones were obtained for LoxPTgFRM2 (clone A and B) and used for phenotypic characterisation.

### 12. Induction of the conditional TgFRM2 KO

To obtain TgFRM2 KO parasites, the loxPTgFRM2-YFP parental line was grown in 50nM rapamycin containing media as described above until fixing. In IFA, TgFRM2 KO parasites were always compared to a control population of untreated loxPTgFRM2-YFP.

To quantify TgFRM2-YFP excision 48h post inoculation, 100 vacuoles were counted for each clone (clone A and B) and condition (RAP treated vs control population). The vacuoles were assessed with regards to their loss of FRM2-YFP signal. A single IFA was counted for each clone. To assess the apicoplast segregation phenotype in FRM2-YFP negative vacuoles, 66 (clone A) and 100 vacuoles (clone B) were counted. For this, quantification was achieved by using the same IFA that was used for the excision rate quantification.

### 13. Transient transfection of CB-EME into *Toxoplasma* parasites

To have parasites transiently expressing CB-EME, 1 × 107 of freshly released TgFRM2-HA or loxPTgFRM2-YFP parasites were transfected with 20 μg DNA by AMAXA electroporation. Subsequently, parasites were grown on HFFs as described above and fixed with 4% paraformaldehyde after 48h or 72h.

## Supporting information

## Author contributions

SD performed *P. falciparum* experiments. JFS performed *T. gondii* experiments. MS produced the TgFRM2-HA and LoxPTgFRM-YFP strains. JW performed bioinformatic analyses. SD and MM conceived the project. SD, MM and JFS wrote the manuscript.

## Acknowledgments

We thank Prof. Mike Blackman for the kind gift of the PKG inhibitor Compound 2 and the B11 DiCre strain. We thank Dr. Jake Baum for the PfACT1 and RON4 antibodies, Dr. Julian Rayner for the MTIP and GAP45 antibodies, Dr Lilach Sheiner for the CPN60 and G2Trx antibodies, Prof Peter Bradley for the Atrx1 antibody and Prof GK Jarori for the enolase antibodies.

---

**Figure S1.**
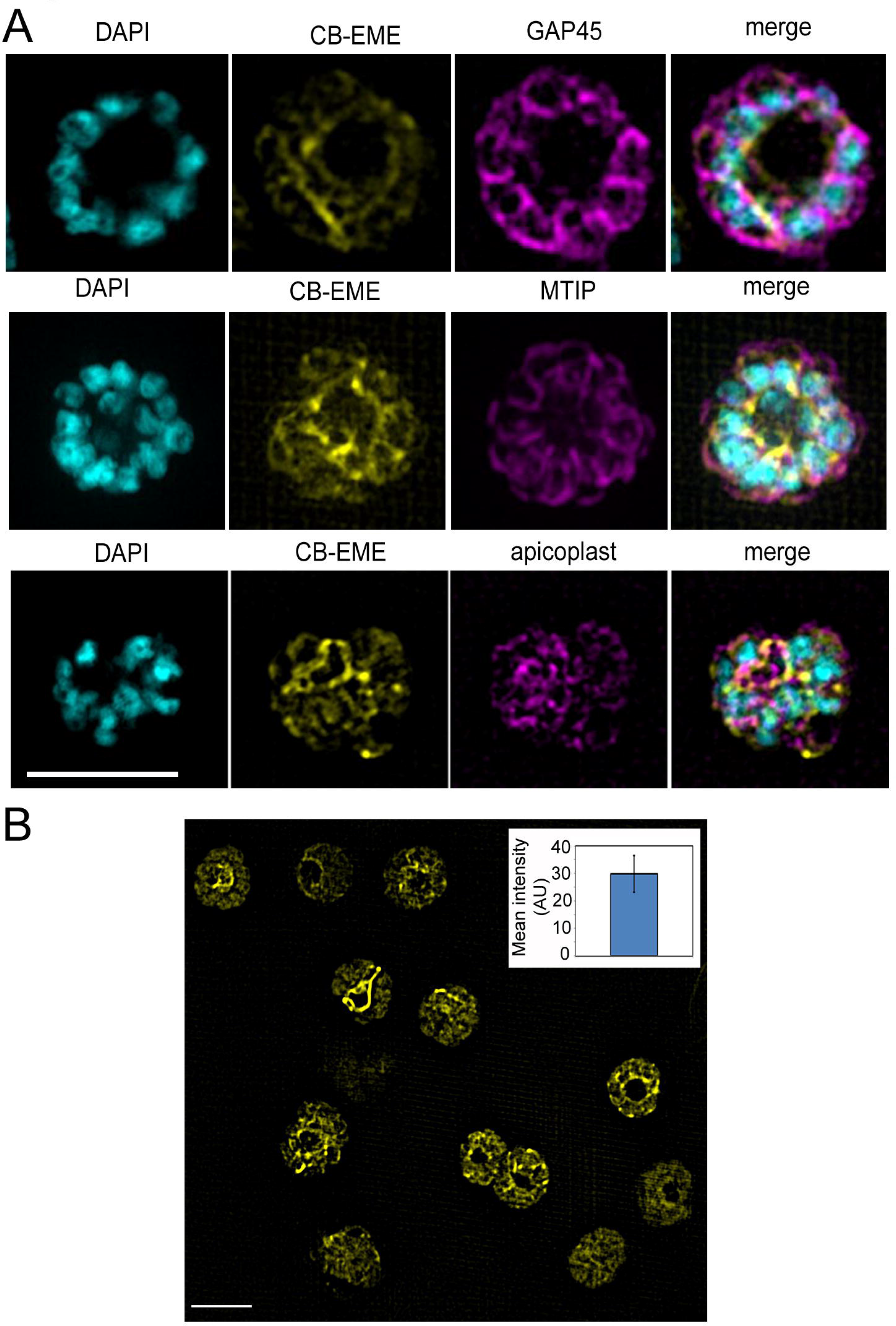
Super-resolution of the CB-EME-labelled F-actin network. **A.** Super-resolution imaging of schizonts show the F-actin network (CB-EME, yellow) co-stained in magenta with IMC-markers GAP45, MTIP and the apicoplast marker CPN60. Note the very close apposition of the apicoplasts with the F-actin network. DAPI stains nuclei (cyan). **B.** Mean fluorescence intensity of the CB-EME channel of each of the schizonts has been presented, N=40. Error bars show SD.

**Figure S2:**
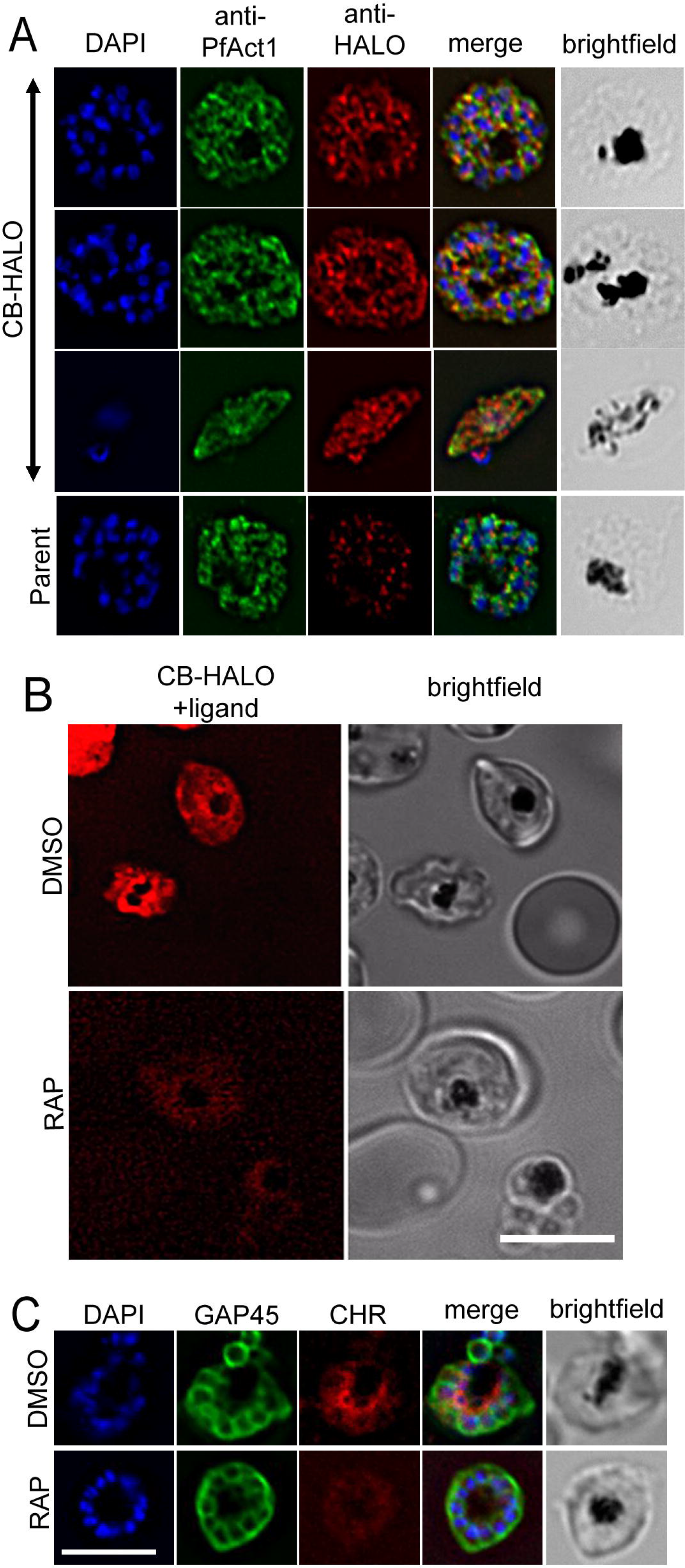
CB-HALO labels the F-actin network similar to CB-EME. **A.** IFA of CB-HALO expressing parasites showing staining with anti-PfACT1 and with an anti-HALO antibody. Note that the anti HALO antibody shows background staining in the parental strain. **B.** Intensity of CB-HALO expression and filament structures (red) are lost upon PfACT1 disruption (RAP). **C.** IFA with anti-GAP45 antibodies show normal formation of merozoites in PfACT1-disrupted parasites (RAP) expressing CB-HALO.

**Figure S3.**
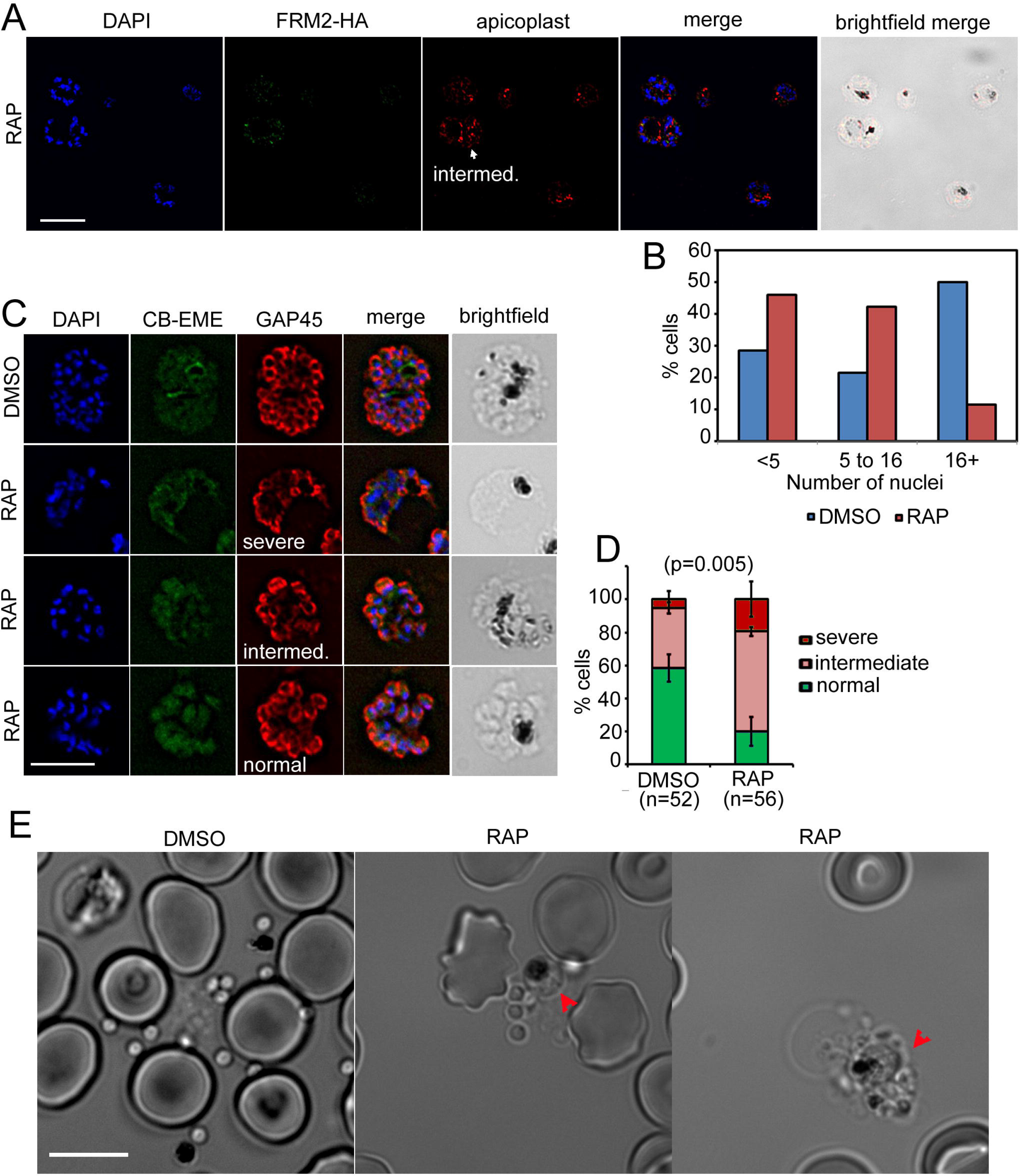
Additional defects in merozoite formation/ cytokinesis in PfFRM2 KO parasites. **A.** IFA image showing schizonts with varying degrees of apicoplast (red) segregation defect 44h post-RAP-treatment of LoxPpffrm2. An example of an intermediate apicoplast defect has been shown (white arrow). Nuclei are stained in blue. **B.** Number of nuclei in the IFA from A were binned to <5, 5 to 16 and 16+ in DMSO controls and RAP-treated parasites. The graph shows a general reduction in the number of nuclei in RAP-treated parasites. **C.** LoxPpffrm2/CBEME parasites were stained with an anti-GAP45 antibody (red) 44h post DMSO/RAP-treatment and examples of the IMC defect have been provided. CBEME fluorescence is shown in green. **D.** Upon quantification of C, normally formed merozoites reduced significantly in PfFRM2 KOs. **E.** When DMSO/RAP-treated schizonts were allowed to egress, conjoined merozoites were apparent in the RAP treated controls (red arrows) but not in DMSO controls.

**Figure S4.**
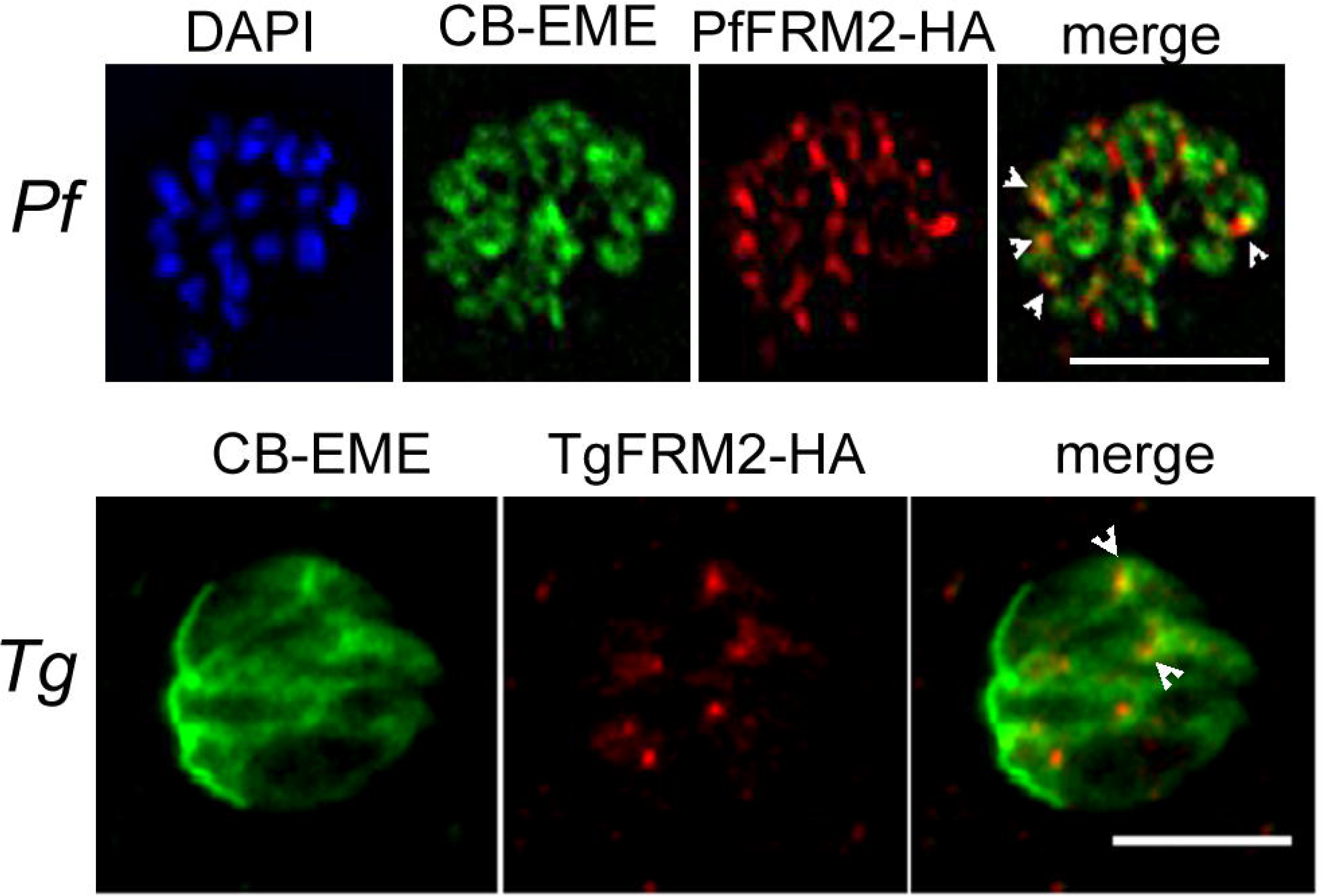
Formin-2 localises to the F-actin network in *P. falciparum* and *T.gondii:* Upper panel. IFA depicting PfFRM2-HA localisation using an anti-HA antibody in *P. falciparum* expressing CB-EME (green). PfFRM2-HA localises to F-actin (merge, white arrows). Nuclei were stained with DAPI (blue). **Lower panel**. IFA showing staining for TgFRM2-HA using an anti-HA antibody in *Toxoplasma* transiently expressing CB-EME (green). TgFRM2-HA localised in close proximity to intracellular actin polymerisation events (merge, white arrows). Parasites were transiently transfected with CB-EME and grown for 48h. Scale bars are 5μm.

**Figure S5.**
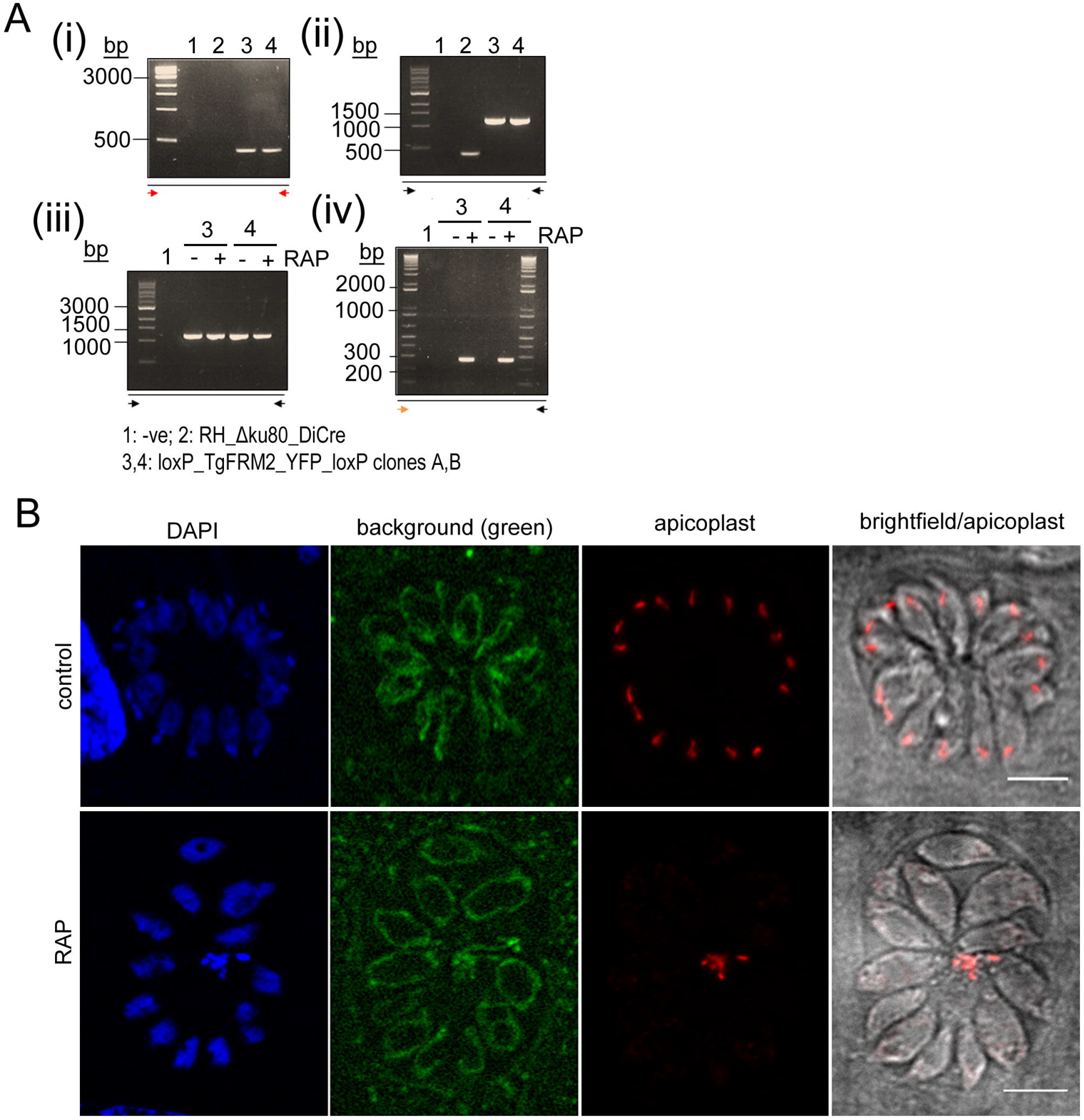
Loss of TgFRM2-YFP upon RAP-treatment. **A.** Integration PCR for TgFRM2-YFP parasites described in **Fig. 7** confirming 5’ integration (i) and 3’ integration (ii). While the amplification of YFP indicates the presence of non-induced parasites in the rapamycin-treated population (iii), excision of TgFRM2-YFP could only be shown for parasites growing under rapamycin (iv). For the excision PCRs (iii, iv) parasites were grown with or without 50nM Rapamycin and mechanically lysed after 48h prior to gDNA collection. Coloured arrows refer to **Fig. 7A** and indicate the sequence amplified by PCR. **B.** IFA depicting apicoplast (anti-CPN60) and background fluorescence in LoxPTgFRM2-YFP parasites (clone B), which outlines the mitochondria. In the control population (without rapamycin treatment), the YFP-tagged FRM2 was not detectable in the absence of an YFP-antibody. RAP-treated parasites show the characteristic mislocalisation of apicoplast material into the residual body, while mitochondria appear unaffected. Parasites were grown for 72h before fixation. Scale bars are 5μm.

**Movie V1, Rapid shape changes of ring stages of *P. falciparum* expressing CB-EME (green)**. Acquisition time is shown in seconds. Scale bar 5μm.

**Movie V2, Dynamic Filopodia-like F-actin extensions from the parasite edges into the RBC cytosol**. Acquisition time is shown in seconds. Scale bar 5μm.

**Movie V3, Dynamic actin filaments in CB-EME expressing parasites (DMSO) are disrupted upon addition of cytochalasin-D**. The green channel shows CB-EME expression. Brightfield images also shown. Acquisition time is shown in seconds. Scale bar 5μm.

**Movie V4, Polar polymerisation of F-actin at the merozoite tip following egress**. Time lapse images of a representative schizont which undergoes egress, followed by polymerisation of F-actin at the merozoite edge (white arrows appearing). Images (green channel, CBEME) and brightfield (greyscale) were acquired every 5 sec. Acquisition time is shown in seconds. Scale bar 5μm.

**Movie V5, F-actin dynamics in gametocytes**. Two representative examples of gametocytes expressing CB-EME show dynamic filaments running along the parasite length and enriched at the tips. Acquisition time is shown in seconds. Scale bar 5μm.

**Movie V6, Multilobular structures of trophozoites are not lost upon addition of cytochalasin-D**.

**Movie V7, CB-EME staining disappears upon conditional genetic deletion of *pfact1***. Ring stage LoxPpfACT1/CBEME parasites were pulse treated with DMSO or RAP for 4h and imaged after 40 hours. CB-EME was imaged in the green channel and shows a disappearance of F-actin upon RAP-treatment. Mitochondria were stained with Mitotracker^®^ (red channel). Acquisition time is shown in seconds. Scale bar 5μm.

**Movie V8, Actin filaments disappear upon genetic deletion of *pffrm2***. Ring stage LoxPpfFRM2/CBEME parasites were DMSO-or RAP-treated for 4h and imaged 40 hours later. CB-EME was imaged in the green channel and shows a disappearance of intracellular F-actin upon RAP-treatment. Acquisition time is shown in seconds. Scale bar 5μm.

## Abbreviations

C2: Compound 2
FV: food vacuole
F-actin: filamentous actin
PfACT1: *P. falciparum* actin-1
PfFRM1: *P. falciparum* Formin-1
PfFRM2: *P. falciparum* Formin-2
TgFRM1: Toxoplasma gondii Formin-1
TgFRM2: Toxoplasma gondii Formin-2
PV: parasitophorous vacuole
RAP: rapamycin
RON4: rhoptry neck protein-4
TJ: tight junction
IMC: inner membrane complex
PCR: Polymerase chain reaction
IFA: (indirect) Immunofluorescence assay
HA: Haemagglutinin
YFP: yellow fluorescent protein
RT: room temperature

